# A sensitive orange fluorescent calcium ion indicator for imaging neural activity

**DOI:** 10.1101/2025.07.28.667269

**Authors:** Abhi Aggarwal, Heather A. Baker, Céline D. Dürst, I-Wen Chen, Pablo de Chambrier, Julia Marie Gonzales, Jonathan S. Marvin, Milène Vandal, Torgny Lundberg, Kenryo Sakoi, Ronak Patel, Ching-Yao Wang, Frank Visser, Yannick Fouad, Smrithi Sunil, Matthew Wiens, Takuya Terai, Kei Takahashi-Yamashiro, Roger J. Thompson, Timothy A. Brown, Yusuke Nasu, Minh Dang Nguyen, Grant R. J. Gordon, Sarah McFarlane, Kaspar Podgorski, Antonius Holtmaat, Robert E. Campbell, Alexander W. Lohman

**Author notes:** Correspondence AWL.

## Abstract

Genetically encoded calcium indicators (GECIs) are vital tools for fluorescence-based visualization of neuronal activity with high spatial and temporal resolution. However, current highest-performance GECIs are predominantly green or red fluorescent, limiting multiplexing options and efficient excitation with fixed-wavelength femtosecond lasers operating at 1030 nm. Here, we introduce OCaMP (also known as O-GECO2), an orange fluorescent GECI engineered from O-GECO1 through targeted substitutions to improve calcium affinity while retaining the favorable photophysical properties of mOrange2. OCaMP exhibits improved two-photon cross-section, responsiveness, photostability, and calcium affinity relative to O-GECO1. In cultured neurons, zebrafish, and mouse cortex, OCaMP outperforms the red GECIs jRCaMP1a and jRGECO1a in sensitivity, kinetics, and signal-to-noise ratio. These properties establish OCaMP as a robust tool for high-fidelity neural imaging optimized for 1030 nm excitation and a compromise-free option within the spectral gap between existing green and red GECIs.

## Introduction

Genetically encoded calcium indicators (GECIs) have revolutionized our ability to visualize neural activity by translating intracellular calcium transients into dynamic fluorescent signals^1–3^. Two-photon (2P) microscopy has become the gold standard for *in vivo* calcium imaging, due to its ability to achieve optical sectioning and deep tissue penetration in brain tissue with minimal photodamage^4^. When paired with GECIs, 2P imaging enables high-speed recording of neural activity in intact brain tissue at the single-cell level.

Traditionally, two-photon excitation has relied on tunable Ti:Sapphire lasers. However, these systems are expensive, power-limited, and require routine maintenance that constrain adoption in many laboratories^4,5^. In contrast, fixed-wavelength lasers, such as ytterbium-doped fiber lasers (YbFLs), offer a promising alternative that operate at wavelengths in the range of 1000-1080 nm, are cost-effective, high-powered, and increasingly integrated into modern imaging platforms^5,6^. However, GECIs that perform optimally under these excitation wavelengths remain limited.

The excitation wavelength of GECIs is closely tied to the fluorescent protein scaffold from which they are derived. GFP-based sensors, such as GCaMP6^7^ and jGCaMP8^8^ provide the highest levels of sensitivity, kinetics, and *in vivo* applicability, but exhibit poor two-photon excitation above 1000 nm due to their declining cross-section beyond ∼950 nm. Red-shifted indicators, typically engineered using mApple-, mRuby-, or mKate-scaffolds, including R-GECO1^9^, jRCaMP1a^10^, and K-GECO^11^ extend excitation beyond 1000 nm but exhibit trade-offs such as poor photostability, slower kinetics, photoswitching, and lysosomal accumulation^12^. Newer variants like mApple-based XCaMP-R^13^ and RCaMP3^14^ have improved response kinetics and dynamic range, but remain sensitive to pH (apparent Ca^2+^-bound p*K*_a_ of 5.3 and 6.1, respectively), and exhibit poor photostability and photoswitching.

Efforts to expand spectral coverage and compatibility at 1030 nm excitation have led to the generation of GECIs based on alternative scaffolds. jYCaMP1^15^ was developed using a yellow fluorescent protein and achieves improved excitation at 1030 nm, but remains less sensitive compared to GCaMP6. More recently, PinkyCaMP^16^, a GECI based on mScarlet has been developed, which offers high sensitivity, photostability, reduced photoswitching, but suffers from slow kinetics. Far-red and near-infrared indicators such as FR-GECO1a^17^ (mKelly2-based), iGECI^18^ (miRFP670 and miRFP720-based), and NIR-GECO variants^19,20^ (miRFP-based) enable two-photon excitation peaks beyond 1100 nm, but do not excite well using 1030 nm fixed-wavelength lasers. While many of the existing GECIs are technically excitable at 1030 nm, none provide an optimal combination of performance and brightness under these imaging conditions.

These trade-offs motivated us to explore mOrange2, an orange fluorescent protein known for its high quantum yield, strong two-photon cross-section near 1030 nm, and exceptional photostability^14,21–23^. These features make it well-suited for applications requiring repeated or prolonged imaging, including high-frame-rate recordings or *in vivo* imaging of dynamic neural activity.

O-GECO1^24^ was the first orange fluorescent calcium indicator engineered from the mOrange2 scaffold^21–23^. It exhibits a large dynamic range (Δ*F*/*F*₀ = 146) and strong two-photon excitation at 1030 nm, but suffers from a relatively low affinity for calcium (*K*_d_ = 1500 nM), limiting its effectiveness for neuronal cytosolic calcium imaging. Given its high sensitivity, 2P cross-section, and photostability, we set out to use O-GECO1 as a starting point for engineering an improved orange GECI optimized for *in vivo* imaging. We hypothesized that incorporating mutations from calcium-binding domains of higher-affinity calcium indicators could yield a new indicator with improved performance while capitalizing on the photophysical properties of mOrange2 and fluorescence change of O-GECO1.

Here, we present the development and characterization of OCaMP (also known as O-GECO2; hereafter referred to as OCaMP), a new orange GECI optimized for *in vivo* imaging. OCaMP exhibits improved calcium affinity while retaining strong fluorescent responses, photostability, and two-photon molecular brightness. We benchmarked OCaMP against jRGECO1a and jRCaMP1a in purified protein, cultured neurons, zebrafish, and performed *in vivo* multiphoton cranial window imaging. Together, these results establish OCaMP as a robust orange GECI with performance suitable for a broad range of experimental modalities.

## Results

### Engineering of OCaMP through rational design guided by high-affinity GECIs

To increase the calcium-binding affinity of O-GECO1 (*K*_d_ = 1500 nM) while retaining its favourable photophysical properties, we performed structure-guided mutagenesis using sequence alignments with high-sensitivity GECIs that exhibit submicromolar calcium affinities, including GCaMP6s (*K*_d_ = 144 nM), jGCaMP8f (*K*_d_ = 334 nM), jRCaMP1a (*K*_d_ = 82 nM), and jRGECO1a (*K*_d_ = 129 nM). Notably, jRGECO1a exhibits the highest sequence similarity to O-GECO1, differing by only 10 amino acids (Supplementary Figure 1), suggesting it could serve as a template for engineering improved binding affinity.

As an initial design, we focused on six of these ten mutations located within the M13 and calmodulin domains, and introduced the corresponding substitutions from jRGECO1a into O-GECO1 (I5N, Q267D, L300F, R339K, V347E, and N358K). However, this construct failed to produce fluorescence when expressed in bacteria. Since Q267 is positioned at the interface between calmodulin and the circularly permuted fluorescent protein, we hypothesized that this residue is critical for proper folding and chromophore maturation. Reverting this substitution (D267Q) restored fluorescence, yielding a variant with five substitutions relative to O-GECO1: I5N, L300F, R339K, V347E, and N358K. We designated this construct as OCaMP (Fig. 1a).

**Figure 1.**
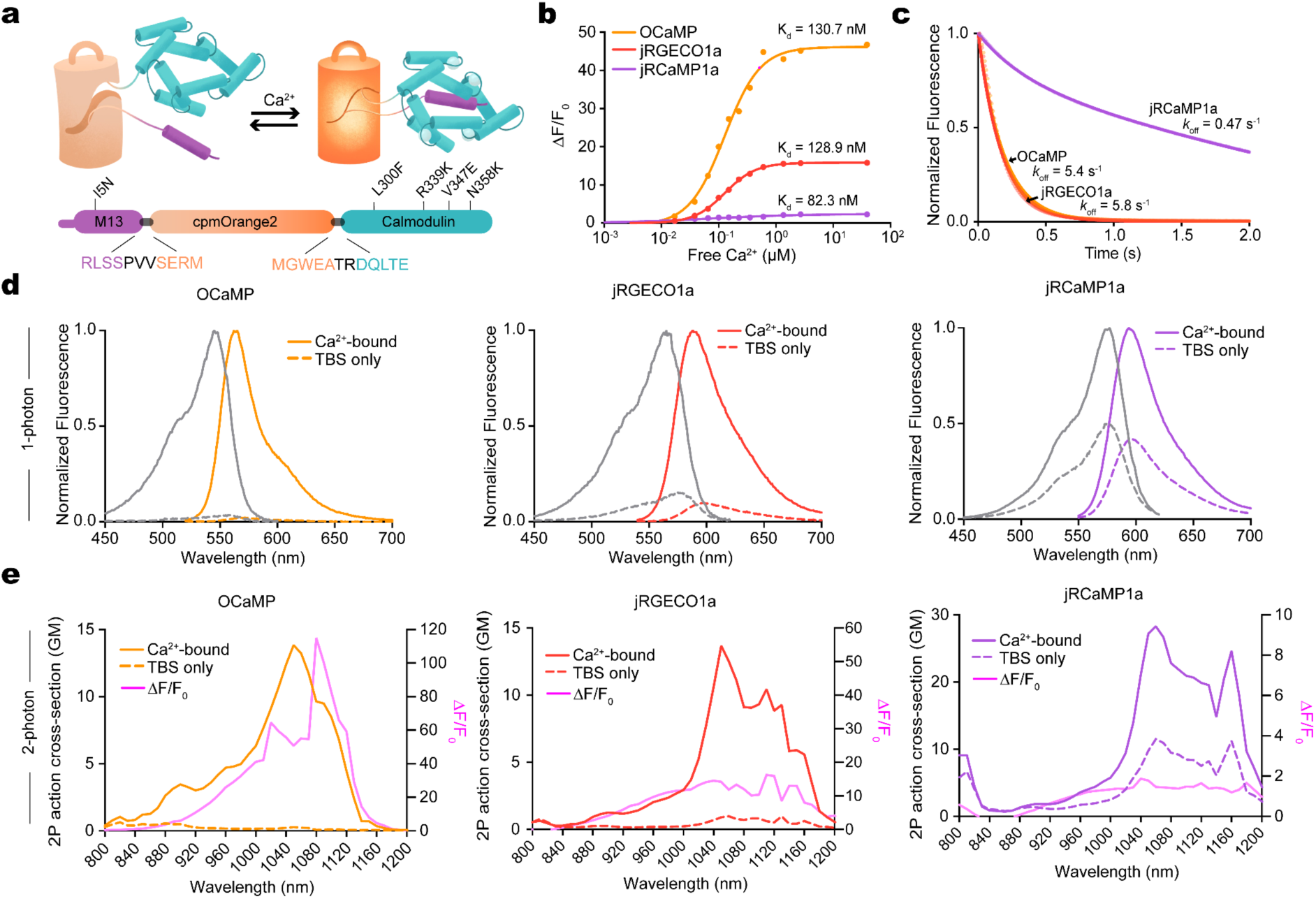
Photophysical properties of OCaMP. (**a**) Protein schematic illustration (top) and linear DNA sequence illustration (bottom). Listed mutations are with respect to O-GECO1. (**b**) Calcium titrations of the indicators as purified protein. (**c**) Stopped-flow measurements of OFF rate of the different indicators as purified protein. (**d**) 1-photon excitation and emission spectra of the indicators in calcium-saturated (10 mM) and calcium-free (TBS) solutions. (**e**) 2-photon excitation spectra and Δ*F*/*F*₀ of the variants in the presence (10 mM) and absence (TBS) of calcium. All measurements were made in triplicates.

In parallel, we tested an alternative design by incorporating the M13-like peptide from jGCaMP8f (residues 1–19, also known as the ENOS peptide^25,26^). The resulting variant, designated OCaMP-fast, exhibited faster decay kinetics but a substantially reduced dynamic range and was not pursued further (Supplementary Fig. 2).

Overall, these experiments guided the construction of OCaMP, a variant of O-GECO1, by incorporating targeted substitutions in the calcium-sensing domains to achieve improved affinity while preserving spectral and structural properties critical for 1030 nm excitation.

### Characterization of OCaMP as purified protein and expressed in mammalian cells

First, we measured the dynamic range and binding affinity of purified OCaMP alongside jRGECO1a and jRCaMP1a (Fig. 1b; see Supplementary Figure 2 for comparison with jYCaMP1, XCaMP-R, and OCaMP-fast). OCaMP displayed a significantly increased calcium affinity (*K*_d_ = 131 nM), more than tenfold higher than O-GECO1, and a large calcium-dependent fluorescence change *in vitro* (ΔF/F₀ = OCaMP: 46.8; jRGECO1a: 11.3; jRCaMP1a: 1.4; Fig. 1b).

OCaMP displayed rapid calcium unbinding kinetics (*k*_off_ = 5.4 s⁻¹), similar to jRGECO1a (5.8 s⁻¹) and substantially faster than jRCaMP1a (0.47 s⁻¹; Fig. 1c). OCaMP’s one-photon excitation and emission peaks are at 545 and 565 nm, respectively, consistent with the mOrange2 fluorescent protein, spectrally separated from jRGECO1a (excitation/emission: 561/589 nm) and jRCaMP1a (excitation/emission: 559/600 nm; Fig. 1d).

In the Ca²⁺-bound state, OCaMP exhibited a quantum yield of 0.22, an extinction coefficient of 62,900 M⁻¹ cm⁻¹, and one-photon molecular brightness of 13.8 (Supplementary Table 1). OCaMP’s quantum yield was similar to jRGECO1a (Φ = 0.20) and lower than jRCaMP1a (Φ = 0.54), and its extinction coefficient exceeded both jRGECO1a (ε = 58,900 M⁻¹ cm⁻¹) and jRCaMP1a (ε = 54,100 M⁻¹ cm⁻¹). Under two-photon excitation, OCaMP exhibited a broad excitation profile from 1000 to 1120 nm and a large peak fluorescence response at 1030 nm (ΔF/F₀ = 42.6), more than 3-fold higher than jRGECO1a (14.1) and over 30-fold higher than jRCaMP1a (1.4; Fig. 1e). In addition, OCaMP showed substantially higher fluorescence responses close to physiological pH, with a Ca²⁺-bound apparent pKₐ of 7.89 than jRGECO1a (5.74) or jRCaMP1a (6.15) (Supplementary Fig. 3).

We validated OCaMP in HeLa cells by applying histamine to evoke intracellular calcium release (Supplementary Fig. 5). OCaMP showed robust fluorescence responses consistent with its high sensitivity observed *in vitro*. To assess light-dependent activation, we exposed cells to brief pulses of 450 nm light under both calcium-free and calcium-bound conditions. As expected, jRGECO1a exhibited substantial fluorescence increases following each 450 nm pulse under both calcium-free and calcium-bound conditions, while jRCaMP1a showed minor increases only in the absence of calcium. In contrast, OCaMP displayed no visible changes in fluorescence during repeated 450 nm exposures in either condition (Supplementary Fig. 4).

We next evaluated photostability under continuous one-photon illumination (exc: 545 nm) in HEK293 cells. Under continuous illumination, OCaMP was substantially more photostable than the other GECIs (fluorescence loss over 120 s: OCaMP, 0.9%; jRGECO1a, 21.0%; jRCaMP1a, 70.4%) (Supplementary Fig. 5)

Together, these results demonstrate that OCaMP combines no photoswitching with high photostability, enabling long-term imaging and reduced optical crosstalk in multiplexed experiments.

### OCaMP shows increased sensitivity to field stimulated neuronal cultures

Next, we expressed OCaMP in cultured rat hippocampal neurons and imaged calcium transients following field stimulation^27^ (Fig. 2a). Neurons expressing OCaMP showed robust cytoplasmic expression and low baseline fluorescence (OCaMP: 18.79 ± 1.75 AU; jRGECO1a: 40.50 ± 3.00 AU; jRCaMP1a: 170.28 ± 9.28 AU; Fig. 2b,f).

**Figure 2.**
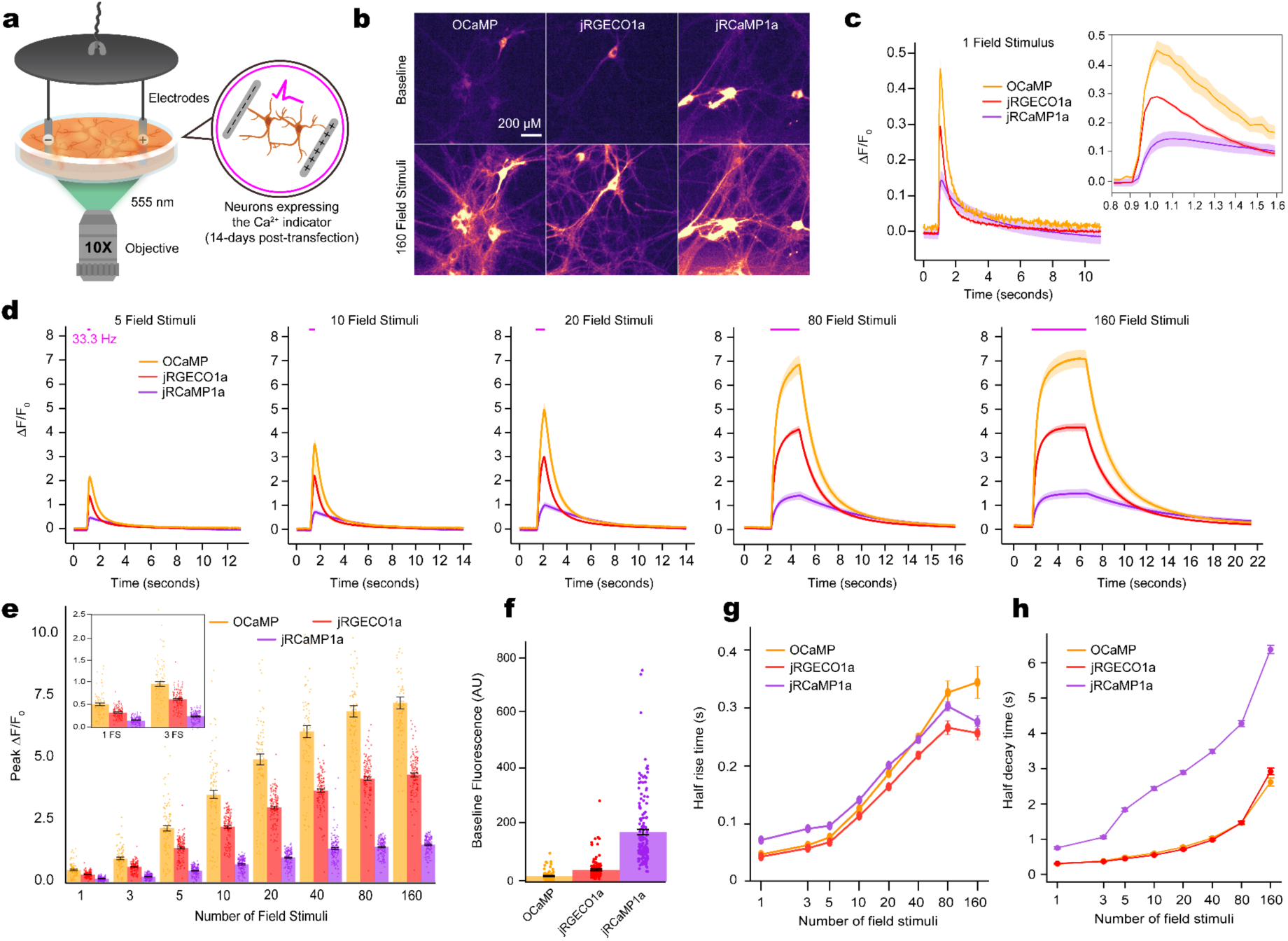
Characterization of OCaMP in cultured neurons. (**a**) Schematic illustration of the primary neuronal culture field stimulation assay. All indicators were excited at 555 nm. (**b**) Baseline fluorescence images of primary rat hippocampal cultured neurons expressing OCaMP, jRGECO1a, and jRCaMP1a. Scale bar denotes 200 µM. (**c**) Δ*F*/*F*₀ traces for the various indicators after applying 1 field stimulus. Right panel is a zoom-in view. (**d**) Average Δ*F*/*F*₀ traces for field stimuli of 5, 10, 20, 80, and 160. (**e**) Peak Δ*F*/*F*₀ for the different indicators across various numbers of field stimuli. (**f**) Baseline brightness of the indicators. (**g**) Half rise time of the indicators across the various number of field stimuli. (**h**) Half decay time. Error bars and shaded regions denote +/- SEM unless otherwise specified.

In response to a single action potential (AP), OCaMP produced a fluorescence response of ΔF/F₀ = 0.49 ± 0.03, which exceeded the responses of jRGECO1a (0.31 ± 0.01) and jRCaMP1a (0.13 ± 0.01; Fig. 2c; see also Supplementary Fig. 2 for comparison with jYCaMP1, XCaMP-R, and OCaMP-fast). Across the full stimulus range (1 to 160 APs), OCaMP consistently had larger peak responses than both red indicators (Fig. 2d–e). For instance, at 160 APs, OCaMP reached a peak ΔF/F₀ of 7.23 ± 0.24, compared to 4.33 ± 0.07 for jRGECO1a and 1.51 ± 0.03 for jRCaMP1a.

OCaMP also exhibited fast kinetics, with half-rise times of 47.36 ± 0.97 ms for OCaMP, 43.42 ± 0.65 ms for jRGECO1a, and 72.11 ± 1.91 ms for jRCaMP1a, and half-decay times of 291.53 ± 9.71 ms, 310.74 ± 7.08 ms, and 751.60 ± 31.38 ms, respectively (Fig. 2g–h).

Together, these results demonstrate that OCaMP offers improved sensitivity, favorable kinetics, and robust performance for detecting neuronal calcium signals compared to existing red-shifted indicators.

### OCaMP shows increased sensitivity to spontaneous neuronal activity in zebrafish larvae

Having demonstrated OCaMP’s performance in cultured neurons, we sought to determine whether it could reliably report spontaneous neuronal activity *in vivo*. We expressed OCaMP, jRGECO1a, or jRCaMP1a in larval zebrafish and performed spontaneous activity imaging using one- and two-photon microscopy (Fig. 3a–j). All indicators labeled broad regions of somata and neuropil (Fig. 3b,f), and OCaMP detected more prominent spontaneous transients compared to jRGECO1a and jRCaMP1a (Fig. 3c,g) under both one- and two-photon excitation.

**Figure 3.**
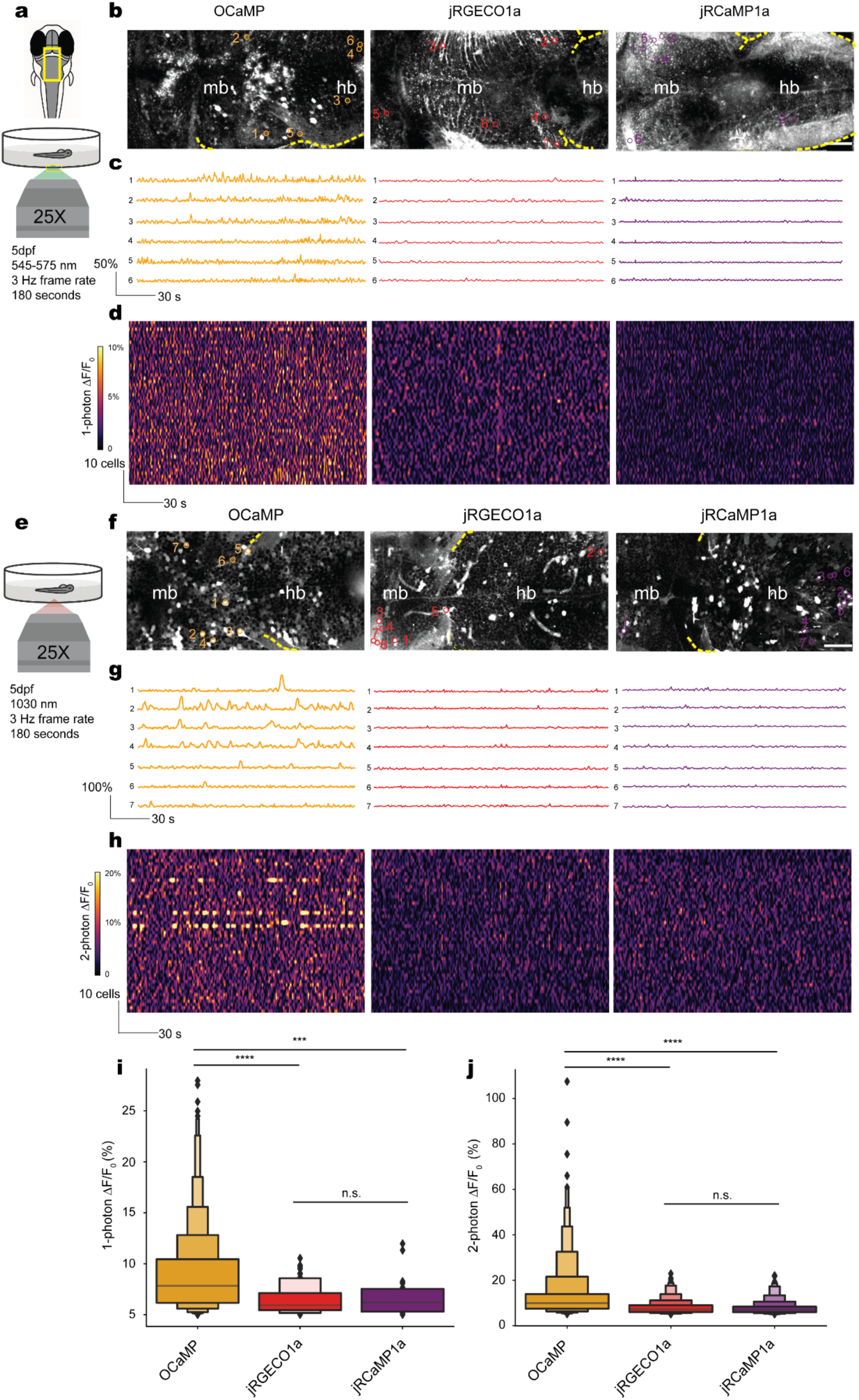
Spontaneous calcium activity imaging in zebrafish larvae using OCaMP, jRGECO1a, and jRCaMP1a. (**a,e**) Experimental schematic showing imaging of spontaneous neural activity in Tol2-injected 5dpf zebrafish larvae expressing OCaMP, jRGECO1a, or jRCaMP1a under (**a**) one-photon (1P, 545 nm for OCaMP, 572 nm for jRCaMP1a, 562 nm for jRGECO1a) and (**e**) two-photon (2P, 1030 nm for all three indicators) excitation. (**b,f**) Maximum intensity projections of the midbrain (mb) and hindbrain (hb) region from representative larvae expressing each indicator imaged with 1P (**b**) or 2P (**f**) microscopy. Yellow lines denote the outline of the fish. Scale bars: 60 μm. (**c,g**) Example ΔF/F₀ traces from selected ROIs in each condition under 1P (**c**) and 2P (**g**) imaging. (**d,h**) Heatmaps of ΔF/F₀ from 50 ROIs per fish, per sensor, under 1P (**d**) and 2P (**h**) conditions. Each row represents one ROI; fluorescence traces are temporally aligned and color-scaled across all panels using a shared lookup table. (**i,j**) Distribution of calcium event amplitudes (ΔF/F₀ peaks) under 1P (**i**) and 2P (**j**) excitation. For 1P imaging, OCaMP (min, Q1, Q2, Q3, max): 0.11, 0.49, 0.73, 1.04, 2.35 (N = 732 ROIs, 6 fish); jRGECO1a: 0.10, 0.22, 0.29, 0.41, 0.93 (N = 59 ROIs); jRCaMP1a: 0.07, 0.18, 0.24, 0.32, 0.61 (N = 27 ROIs). For 2P imaging, OCaMP: 0.09, 0.42, 0.60, 0.83, 2.10 (N = 569 ROIs); jRGECO1a: 0.12, 0.31, 0.47, 0.65, 1.75 (N = 377 ROIs); jRCaMP1a: 0.06, 0.20, 0.27, 0.36, 1.14 (N = 349 ROIs). Boxplots show the median (horizontal line), interquartile range (box), and individual data points, with whiskers extending to Q1 − 1.5×IQR and Q3 + 1.5×IQR. Kruskal–Wallis test with Dunn’s multiple comparisons; ****P < 0.0001; n.s. = non-significant

Heatmaps of ΔF/F₀ across the top 50 ROIs showed consistently stronger signals for OCaMP across both excitations (Fig. 3d,h). Median ΔF/F₀ values reached 7.83% and 9.91% for OCaMP under one- and two-photon excitation, respectively, compared to jRGECO1a (5.95%, 7.06%) and jRCaMP1a (6.21%, 6.88%) (Fig. 3i–j).

These results highlight OCaMP’s increased sensitivity for detecting spontaneous neuronal activity in the intact zebrafish brain across both one- and two-photon imaging.

### *In vivo* two-photon imaging of spontaneous and sensory-evoked activity in mouse cortex

To assess OCaMP’s utility for *in vivo* functional imaging, we expressed OCaMP, jRGECO1a, and jRCaMP1a in layer 2/3 pyramidal neurons of the mouse barrel cortex and performed two-photon imaging in awake, head-fixed mice (Fig. 4a–b). OCaMP-expressing neurons exhibited larger spontaneous transients than those expressing the red indicators (Fig. 4c).

**Figure 4.**
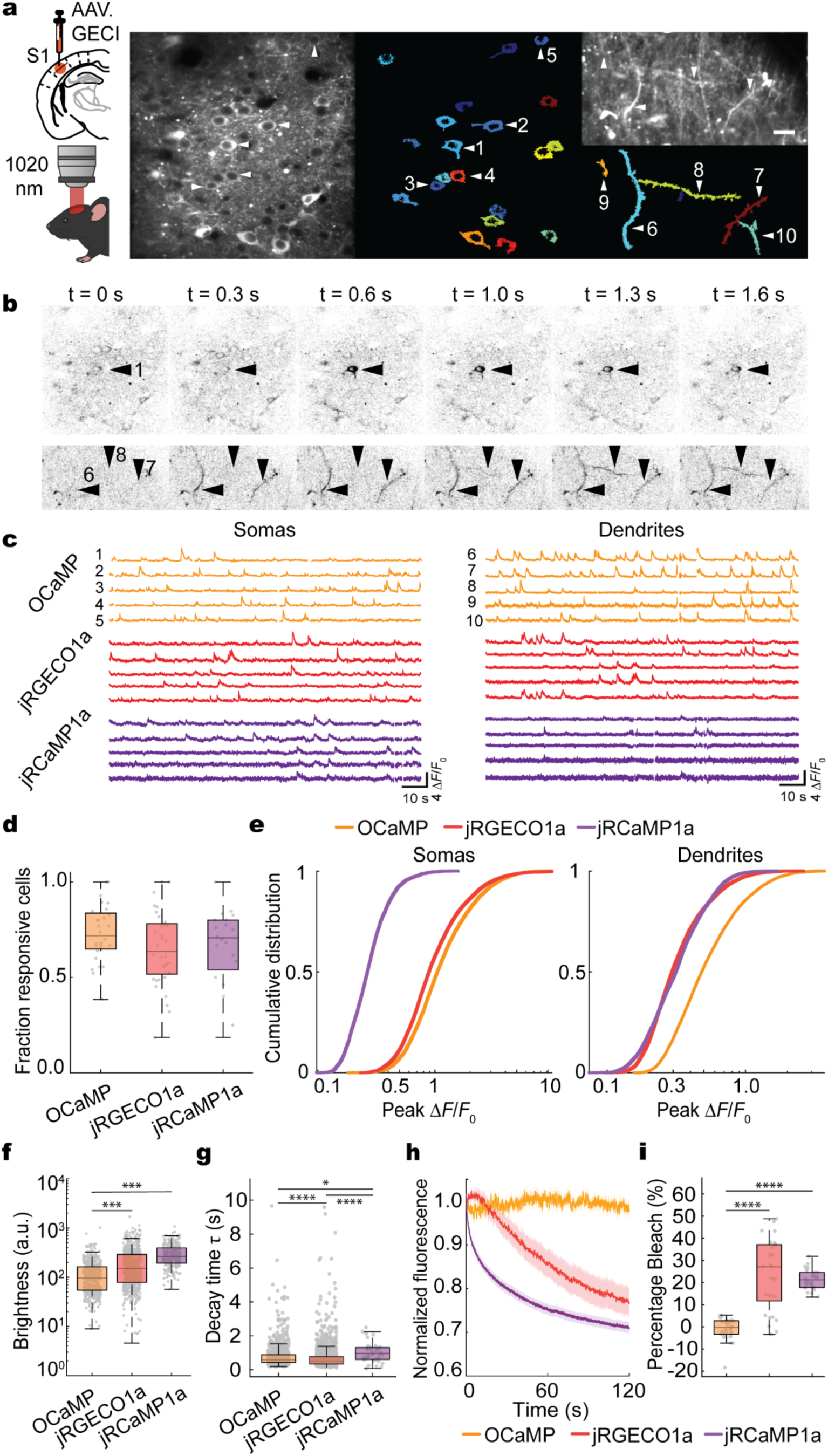
*In vivo* imaging of spontaneous calcium activity in mouse layer 2/3 barrel cortex neurons. (**a**) Experimental scheme with an example of single-plane average projection image of the motion-corrected time series of OCaMP expressing neurons and extracted ROIs from somas and dendrites or axons. (**b**) Time-lapses of the examples shown in (**a**). (**c**) Example traces of OCaMP from the example shown in (**a**), jRGECO1a, and jRCaMP1a. (**d**) Fraction of cells showing at least two responses during 2 min of recording for OCaMP (min, Q1, Q2, Q3 and max), 0.38, 0.65, 0.72, 0.84, 1.00, (*n* = 28 field of views (FOVs) from 4 mice); for jRGECO1a (min, Q1, Q2, Q3 and max) 0.19, 0.52, 0.64, 0.78, 1.00, (*n* = 37 FOVs from 6 mice); and for jRCaMP1a (min, Q1, Q2, Q3 and max) 0.19, 0.54, 0.71, 0.80, 1.00 (*n* = 20 FOVs from 3 mice). Data passed Shapiro-Wilk normality test (α = 0.05 level). One-way ANOVA test: *P* = 0.22. (**e**) *Left*: Distribution of Ca^2+^ events amplitudes (Δ*F*/*F*0) measured in somas of L2/3 cells for OCaMP (min, Q1, Q2, Q3, max in Δ*F*/*F*0) 0.18, 0.73, 1.05, 1.68, 22.99 (*n* = 494, 28 FOVs, 4 mice), jRGECO1a (min, Q1, Q2, Q3, max in Δ*F*/*F*0) 0.23, 0.64, 0.89,1.40, 36.69 (*n* = 643 cells, 37 FOVs, 6 mice), and jRCaMP1a (min, Q1, Q2, Q3, max in Δ*F*/*F*0) 0.10, 0.20, 0.27, 0.34, 1.58 (*n* = 246 cells, 20 FOVs, 3 mice). Kruskal–Wallis test *P* < 0.0001. Dunn’s multiple comparisons test: OCaMP vs. jRGECO1a: *P* < 0.0001, OCaMP vs. jRCaMP1a: *P* < 0.0001, jRGECO1a vs. jRCaMP1a: *P* < 0.0001. *Right:* Distribution of response amplitudes (Δ*F*/*F*0) measured in dendrites and axons of cortical L1 for OCaMP (min, Q1, Q2, Q3, max in Δ*F*/*F*0) 0.16, 0.34, 0.48, 0.76, 3.71 (*n* = 765 ROIs, 42 FOVs, 5 mice), jRGECO1a (min, Q1, Q2, Q3, max in Δ*F*/*F*0) 0.10, 0.22, 0.29, 0.43, 2.63 (*n* = 556 ROIs, 34 FOVs, 6 mice), and jRCaMP1a (min, Q1, Q2, Q3, max in Δ*F*/*F*0) 0.07, 0.22, 0.31, 0.45, 1.73 (*n* = 368 ROIs, 26 FOVs, 4 mice). Kruskal–Wallis test *P* < 0.0001. Dunn’s multiple comparisons test: OCaMP vs. jRGECO1a: *P* < 0.0001, OCaMP vs. jRCaMP1a: *P* < 0.0001, jRGECO1a vs. jRCaMP1a *P* > 0.999. (**f**) Baseline brightness measured as the 30^th^ percentile fluorescence (during 2 min of recording) for OCaMP (min, Q1, Q2, Q3 and max in a.u.): 8.9, 54.6, 95.7, 162.1, 1019.3 (*n* = 494 cells, 28 FOVs, 4 mice), jRGECO1a (min, Q1, Q2, Q3 and max in a.u.): 4.6, 78.2, 150.6, 290.7, 1691.9 (*n* = 644 cells, 37 FOVs, 6 mice) and jRCaMP1a (min, Q1, Q2, Q3 and max in a.u.): 56.6, 192.8, 263.6, 393.6, 1072.5 (*n* = 246 cells, 20 FOVs, 3 mice). Kruskal-Wallis test; *P* < 0.0001. Dunn’s multiple comparisons test with OCaMP as a control: OCaMP versus jRGECO1a: *P* < 0.001, OCaMP versus jRCaMP1a: *P* < 0.001. (**g**) Decay time constants (τ) measurements of somatic spontaneous detected responses for OCaMP (min, Q1, Q2, Q3, max in sec): 0.19, 0.44, 0.60, 0.89, 9.66 (*n* = 494, 28 FOVs, 4 mice), jRGECO1a (min, Q1, Q2, Q3, max in sec): 0.12, 0.35, 0.51, 0.77, 9.58 (*n* = 643 cells, 37 FOVs, 6 mice), and jRCaMP1a (min, Q1, Q2, Q3, max in sec): 0.08, 0.61, 0.97, 1.32, 2.5 (*n* = 246 cells, 20 FOVs, 3 mice). Kruskal– Wallis test *P* < 0.0001. Dunn’s multiple comparisons test: OCaMP vs. jRGECO1a: *P* < 0.0001, OCaMP vs. jRCaMP1a: *P* = 0.0078, jRGECO1a vs. jRCaMP1a *P* <0.0001. (**h**) Average traces with shaded areas representing averages +/- SEM of the brightness normalized to their initial value (t = 0 to t=8333 ms) for OCaMP (orange), jRGECO1a (red), and jRCaMP1a (purple). (**i**) Bleach percentage is measured as the percentage change from the mean intensity over the first 10 s until the last 10 s of each 120 s long recording for OCaMP (median change: -0.2 %, IQR = 5.9 %), jRGECO1a (median change: 27.0%, IQR = 25.3%,), and jRCaMP1a (median change: 21%, IQR = 6.7%,). Kruskal–Wallis test *P* < 0.0001. Dunn’s multiple comparisons test: OCaMP vs. jRGECO1a: *P* < 0.0001, OCaMP vs. jRCaMP1a: *P* < 0.0001, jRGECO1a vs. jRCaMP1a *P* > 0.999. Data were acquired from *n* = 38, FOVs, 4 mice for OCaMP, *n* = 29 FOVs (4 mice) for jRGECO1a, and *n* = 28 FOVs (5 mice) for jRCaMP1a. Boxplots display the median (horizontal line), interquartile range (box), and individual data points with the lower and upper whiskers representing Q1-1.5*IQR and Q3+1.5*IQR, respectively. Statistical significance is indicated as follows: *P* < 0.05 (*), *P* < 0.01 (**), *P* < 0.001 (**\*\*\***), and *P* < 0.0001 (****). Comparisons without asterisks are not statistically significant.

The proportion of somatic cells exhibiting at least two spontaneous calcium transients during a 2-minute recording was comparable across indicators (median (interquartile range (IQR)): OCaMP: 0.72 (0.19), jRGECO1a: 0.64 (0.26), jRCaMP1a: 0.71 (0.26) Fig. 4d). The amplitude of somatic events was substantially larger with OCaMP (ΔF/F₀ median (IQR) = 1.05 (0.95)) than with jRGECO1a (0.89 (0.76)) or jRCaMP1a (0.27 (0.14)) (Fig. 4e, left). In dendrites, OCaMP also produced higher median (IQR) response amplitudes (0.48 (0.42)) compared to jRGECO1a (0.29 (0.21)) and jRCaMP1a (0.31 (0.23)) (Fig. 4e, right).

During whisker stimulation, OCaMP-expressing neurons exhibited clear, time-locked calcium responses with peak amplitudes that exceeded those of the red indicators (2.48 ± 0.19 for OCaMP, 1.00 ± 0.08 for jRGECO1a, and 0.89 ± 0.10 for jRCaMP1a; Supplementary Fig. 6).

Baseline fluorescence intensity was lower in OCaMP-expressing neurons (median (IQR) 95.7 a.u. (107.5)) compared to jRGECO1a (150.6 a.u. (212.5)) and jRCaMP1a (263.6 a.u. (200.8)) (Fig. 4f). Spontaneous decay time constants were median (IQR) 0.60 s (0.45) for OCaMP, 0.51 s (0.42) for jRGECO1a, and 0.97 s (0.71) for jRCaMP1a (Fig. 4g). During continuous imaging, OCaMP showed minimal photobleaching (median (IQR): −0.2% (5.9%), whereas jRGECO1a and jRCaMP1a exhibited larger signal losses (−27.0% (25.3%) and −21.0% (6.7%), respectively) (Fig. 4h,i).

Together, these results establish OCaMP as a robust, photostable, and sensitive calcium indicator for *in vivo* imaging, capable of resolving both spontaneous and sensory-evoked activity with higher signal amplitude and photostability than existing red-shifted sensors.

### OCaMP enables reliable detection of single action potentials *in vivo*

Next, we evaluated OCaMP’s performance *in vivo* by performing simultaneous two-photon calcium imaging and juxtacellular recordings in layer 2/3 neurons of the mouse barrel cortex (Fig. 5a). Fluorescence signals were recorded from OCaMP- or jRGECO1a-expressing cells during spontaneous activity.

**Figure 5.**
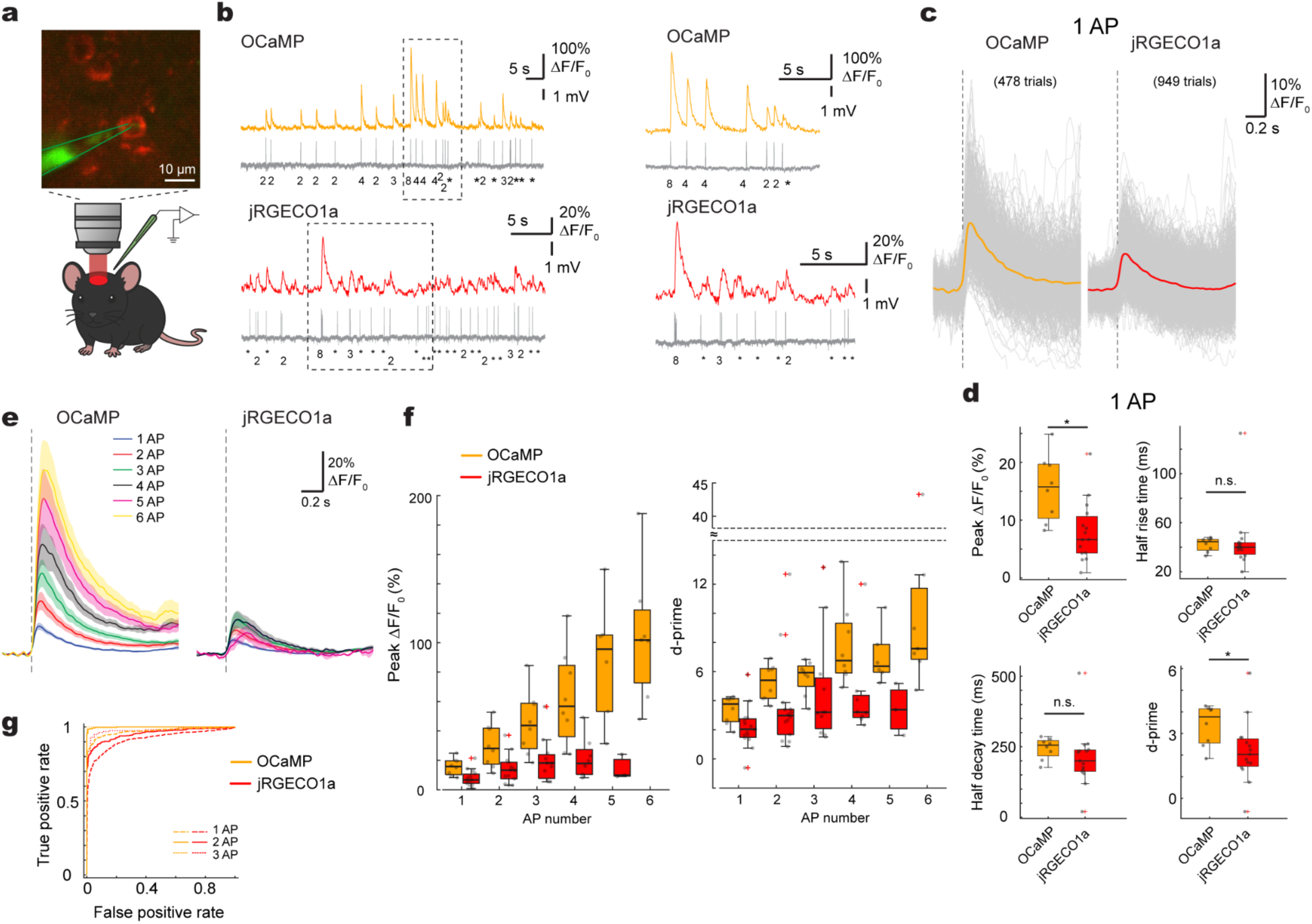
OCaMP enables reliable detection of single action potentials *in vivo*. (a) Experimental setup. Top, example image of an OCaMP-expressing neuron in layer 2/3 of mouse barrel cortex. Bottom, schematic of the in vivo imaging and stimulation configuration in the mouse. Scale bar = 10µm. (b) Simultaneous fluorescence and electrophysiological recordings from representative neurons expressing OCaMP (top) or jRGECO1a (bottom). Action potentials (AP) were of spontaneous activity.The AP numbers of spiking events were indicated below the traces. Asterisks indicated spiking events of single AP. Right, zoomed-in views of traces corresponding to the dashed boxes at the left. (c) Fluorescence responses aligned to single AP’s peak latencies (vertical dashed lines) from all representative neurons (478 trials from 9 cells for OCaMP, and 949 trials from 17 cells for jRGECO1a). Gray lines represent individual trials; colored trace shows mean ± s.e.m. Left, OCaMP; right, jRGECO1a. (d) Quantification of single AP responses. Box plots show peak ΔF/F₀, half rise time (t½ rise), half decay time (t½ decay), and d-prime for neurons expressing OCaMP or jRGECO1a. Statistical comparisons by 2-sample t-test; *n* = 8 OCaMP cells from 3 mice, 15 jRGECO1a cells from 4 mice. p = 0.0044, 0.77, 0.39, 0.047 for comparisons of peak ΔF/F₀, half rise time, half decay time, and d-prime respectively. Asterisks indicated p<0.05. n.s. = non-significant. (e) Grand averaged fluorescence responses across average traces of individual cells to increasing numbers of evoked AP for neurons expressing OCaMP (left; n=8, 8, 8, 8, 6, 7 cells for 1-6 AP) or jRGECO1a (right; n=15, 14, 11, 8, 3 cells for 1-5 AP). Each trace represents the mean response (± s.e.m.) across trials for the indicated spike count. (f) Quantification of peak ΔF/F₀ and d-prime across increasing numbers of evoked action potentials (1–6). Box plots compare responses from neurons expressing OCaMP and jRGECO1a. Box-plot for jRGECO1a at AP = 6 was omitted due to n < 3. Statistical comparisons between AP numbers by 1-way ANOVA: p<0.001, <0.001, p=0.015, 0.040 for peak ΔF/F₀, half rise time, half decay time, and d-prime respectively for OCaMP; p=0.036, 0.0015, 0.44, 0.23 respectively for jRGECO1a. (g) ROC curves for detecting 1, 2, and 3 action potentials based on fluorescence responses. OCaMP achieves higher detection accuracy than jRGECO1a across all spike counts tested. AUC: 0.971 (OCaMP; n=478 trials) vs 0.918 (jRGECO1a; n=949 trials) for 1 AP; 0.999 (OCaMP; n=274 trials) vs 0.955 (jRGECO1a; n=430 trials) for 2 AP; 1.000 (OCaMP; n=148 trials) vs 0.978 (jRGECO1a; n=159 trials) for 3 AP.

Fluorescence signals were time-locked to single action potentials and revealed clear calcium transients in both indicators (Fig. 5b). In response to single spikes, OCaMP generated larger fluorescence changes than jRGECO1a (Fig. 5c). Peak ΔF/F₀ values for single APs were significantly higher in OCaMP-expressing neurons (15.58 ± 2.04%, n=8 cells) than in jRGECO1a (7.98 ± 1.36%, n=15 cells; *p* = 0.0044, 2-sample t-test; Fig. 5d). Response kinetics were comparable, with half rise times of 42.25 ± 1.98 ms (OCaMP) and 44.93 ± 6.59 ms (jRGECO1a) (*p* = 0.77, 2-sample t-test), and half decay times of 244.13 ± 13.42 ms and 210.40 ± 26.83 ms, respectively (*p* = 0.39, 2-sample t-test). d-prime values for single-spike detection were also significantly greater for OCaMP (3.38±0.33) than jRGECO1a (2.18±0.37) (*p* = 0.047, 2-sample t-test; Fig. 5d).

To assess performance across a range of spike counts, we recorded fluorescence responses to spiking events of different numbers of action potentials. OCaMP showed graded increases in ΔF/F₀ with spike number and consistently outperformed jRGECO1a in both amplitude and d-prime across all tested conditions (Fig. 5e,f). Notably, the difference in ΔF/F₀ became more pronounced at higher spike counts (2–6 APs), suggesting that OCaMP exhibits a broader dynamic range and reduced saturation compared to jRGECO1a. ROC analysis further confirmed improved spike detectability with OCaMP, yielding higher AUC values for 1, 2, and 3 action potentials (Fig. 5g; AUC = 0.971 vs 0.918 for OCaMP vs jRGECO1a for 1 AP, 0.999 vs 0.955 for 2 AP, 1.000 vs 0.978 for 3 AP).

These findings demonstrate that OCaMP enables more sensitive detection of single- and multi-spike activity *in vivo* compared to jRGECO1a, with increased fluorescence amplitude and discriminability, and similar kinetics.

## Discussion

Here, we describe OCaMP, an orange genetically encoded calcium indicator optimized for high performance under two-photon excitation at 1030 nm. Starting from the O-GECO1 scaffold, we introduced targeted substitutions from jRGECO1a to increase calcium affinity while preserving the favorable photophysical properties of mOrange2. The resulting sensor is bright, photostable, and sensitive, with broad utility for deep-brain imaging using fixed-wavelength industrial lasers.

Relative to O-GECO1, OCaMP exhibits a >10-fold improvement in calcium affinity (*K*_d_ = 130 nM vs. 1,500 nM), enabling detection of single action potentials *in vivo*. Across cultured neurons, zebrafish, and mouse cortex, OCaMP consistently outperformed jRGECO1a and jRCaMP1a in response amplitude, dynamic range, and signal-to-noise ratio. Its expression time was also notably shorter (∼3–6 weeks), offering practical advantages for longitudinal studies.

OCaMP retained fast kinetics and demonstrated robust photostability under continuous illumination. Unlike jRGECO1a, it showed no detectable photoactivation when exposed to repeated 450 nm pulses in either calcium-bound or calcium-free conditions. This resistance to light-induced activation, combined with minimal photobleaching, makes OCaMP particularly well-suited for experiments involving blue-light stimulation or multi-color imaging for long durations.

Although mOrange2 has been underused in sensor development, our findings suggest it is a promising scaffold for future GECI engineering. OCaMP’s emission profile is well-separated from existing green and red indicators, supporting potential multiplexed imaging applications. Further work will be needed to evaluate performance in deeper brain structures and in combination with optical actuators.

Taken together, these results position OCaMP as a strong addition to the GECI toolkit. Its high sensitivity, fast kinetics, photostability, and compatibility with 1030 nm excitation address key limitations of current red-shifted indicators, making it a compromise-free option for two-photon and multi-color imaging.

---

**Supplementary Table 1.** Photophysical properties of OCaMP and other red-shifted GECIs. Properties for OCaMP, jRGECO1a, and jRCaMP1a were measured in parallel under identical conditions. ^a^Properties for O-GECO1 were previously reported. ^b^2P Δ*F*/*F*₀ values for O-GECO1 were measured in HeLa cells.

| Variant Name | OCaMP |  | jRGECO1a |  | jRCaMP1a |  | O-GECO1<sup>a</sup> |  |
| --- | --- | --- | --- | --- | --- | --- | --- | --- |
|  | APO | SAT | APO | SAT | APO | SAT | APO | SAT |
| 1-photon excitation maxima $\lambda_{exc}$ (nm) | 545 | | 565 | | 577 | | 545 | |
| 1-photon emission maxima $\lambda_{exc}$ (nm) | 565 | | 588 | | 594 | | 565 | |
| 1-photon $\Delta F/F_0$ | 46.8 | | 11.3 | | 1.4 | | 146 | |
| $K_d$ (nM) | 131 | | 129 | | 82 | | 1500 | |
| Apparent Hill coefficient ($n_H$) | 0.63 | | 1.6 | | 0.52 | | 2.06 | |
| $k_{off}$ (s<sup>-1</sup>)<br>($t_{1/2}$ in parentheses) | 5.4 (184 ms) | | 5.8 (172 ms) | | 0.47 (2.2 s) | | 6.2 (161 ms) | |
| Apparent pKa | 9.42 | 7.89 | 8.74 | 5.74 | 5.64 | 6.15 | 9.52 | 6.24 |
| Extinction Coefficient $\epsilon$ (M<sup>-1</sup>cm<sup>-1</sup> × 1000) | 3.6 | 62.9 | 11.2 | 58.9 | 33.8 | 54.1 | 2.0 | 62.9 |
| Quantum yield $\Phi$ | 0.11 | 0.22 | 0.13 | 0.20 | 0.29 | 0.54 | 0.07 | 0.22 |
| 1-Photon Brightness (×1000) | 0.40 | 13.8 | 1.5 | 11.8 | 9.8 | 29.2 | 0.14 | 12.0 |
| 2-Photon $\Delta F/F_0$, (1030 nm) | 42.6 | | 14.1 | | 1.4 | | 41.0<sup>b</sup> | |

**Supplementary Figure 1.**
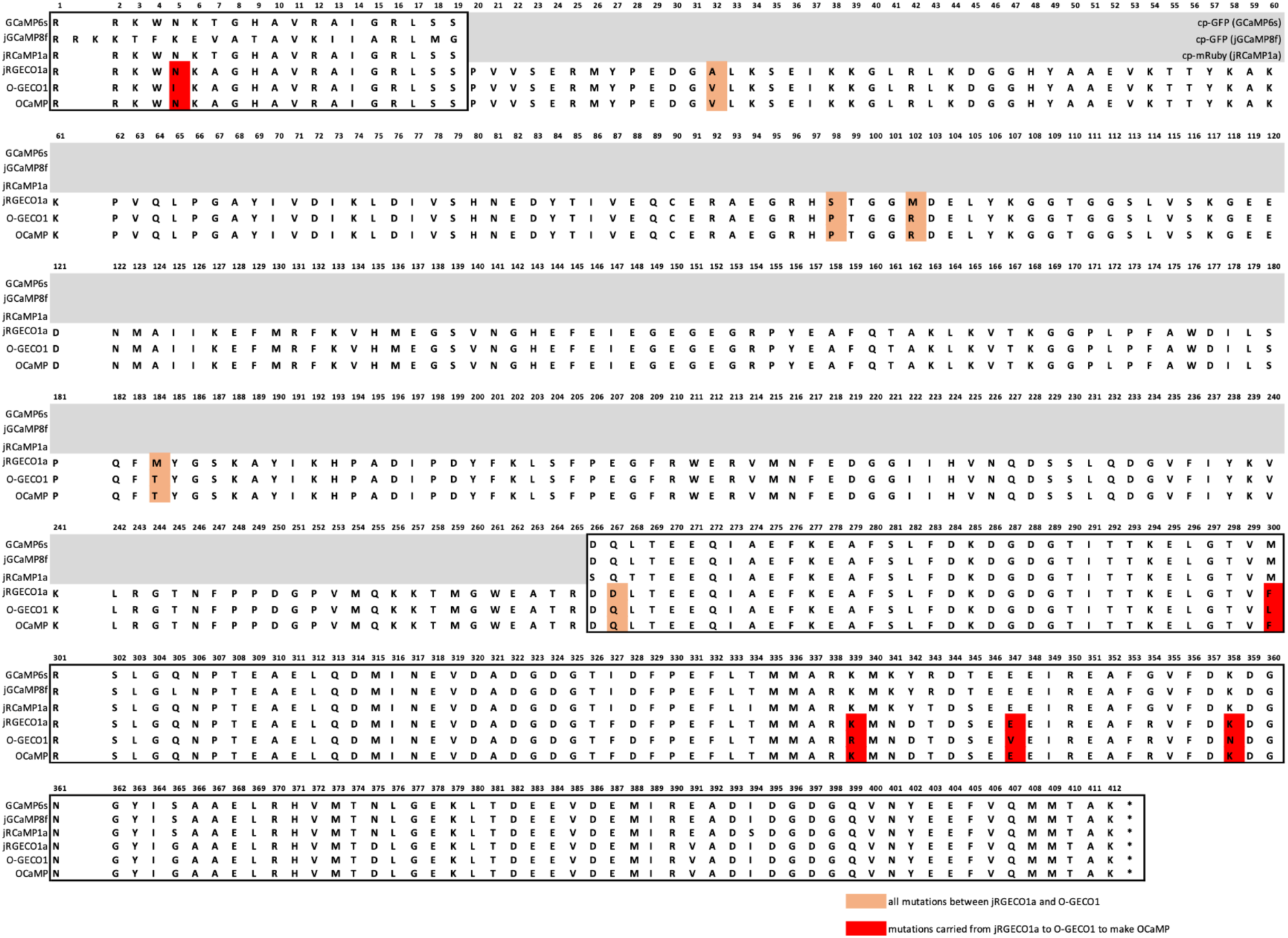
Sequence alignment of OCaMP and related GECIs. Amino acid alignment of OCaMP, O-GECO1, jRGECO1a, jRCAMP1a, jGCaMP8f and GCaMP6s. Boxed regions indicate the M13 peptide (residues 1-19) and calmodulin domain (residues 266-412). All residues that differ between jRGECO1a and O-GECO1 are shaded in beige or red. Residues transferred from jRGECO1a to O-GECO1 to develop OCaMP are highlighted in red. Grayed sequences containing circularly permuted fluorescent proteins with lower sequence identity to OCaMP are omitted for clarity.

**Supplementary Figure 2.**
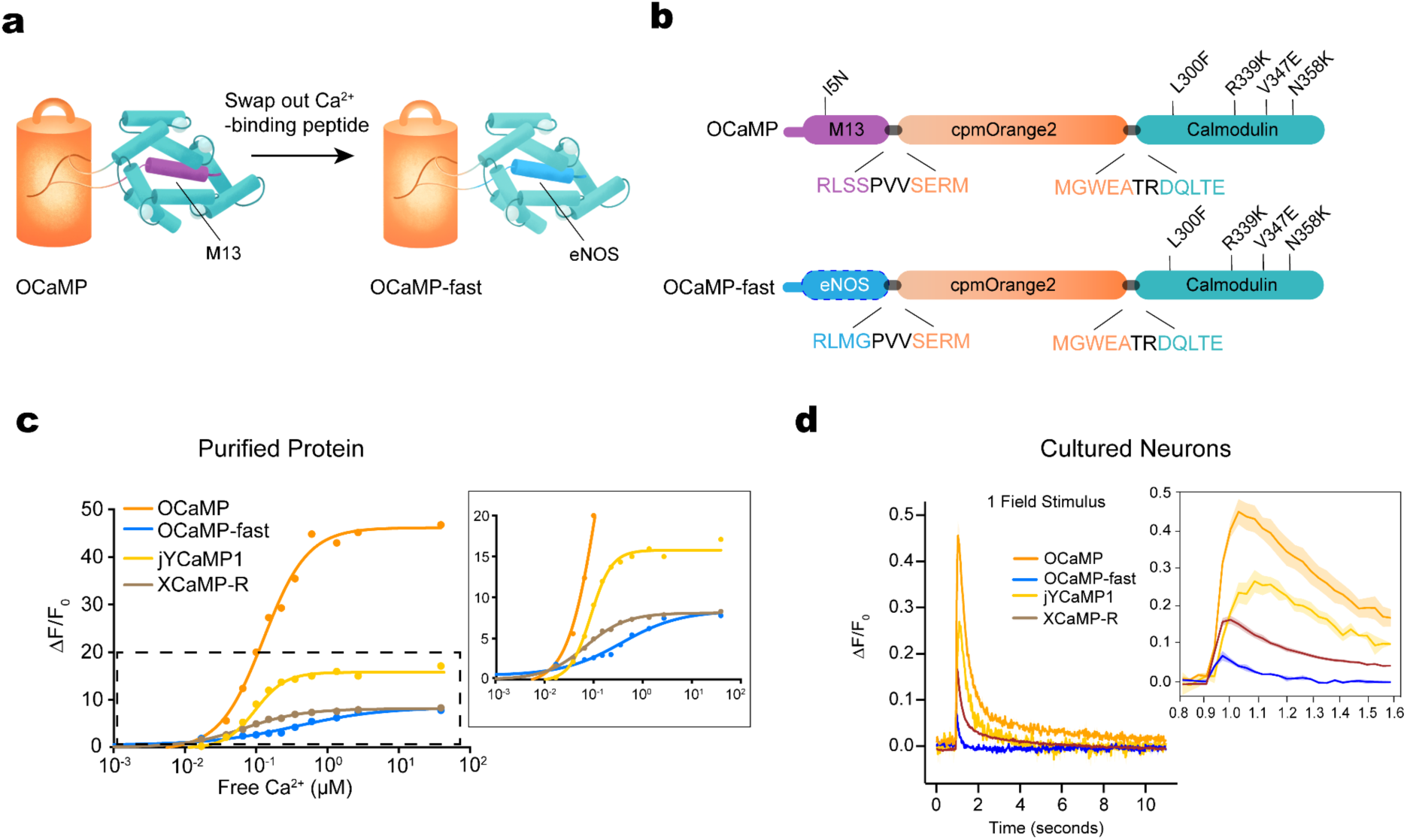
Comparison of OCaMP with OCaMP-fast, XCaMP-R, and jYCaMP1 in purified protein and cultured neurons. (**a**) Protein representation and (**b)** sequence schematic of OCaMP and OCaMP-fast. OCaMP-fast incorporates the eNOS peptide from jGCaMPf in place of the original M13 domain. (**b**) Calcium titration curves of purified protein for OCaMP (orange), OCaMP-fast (blue), jYCaMP1 (yellow), and XCaMP-R (brown). (**c**) Fluorescence responses in cultured neurons following a single field stimulus. OCaMP-fast exhibited faster decay kinetics but reduced ΔF/F₀ compared to OCaMP. jYCaMP1 and XCaMP-R are included for reference. Inset shows expanded view from 0.8 to 1.8 seconds post-stimulus. Shaded areas represent mean ± SEM.

**Supplementary Figure 3.**
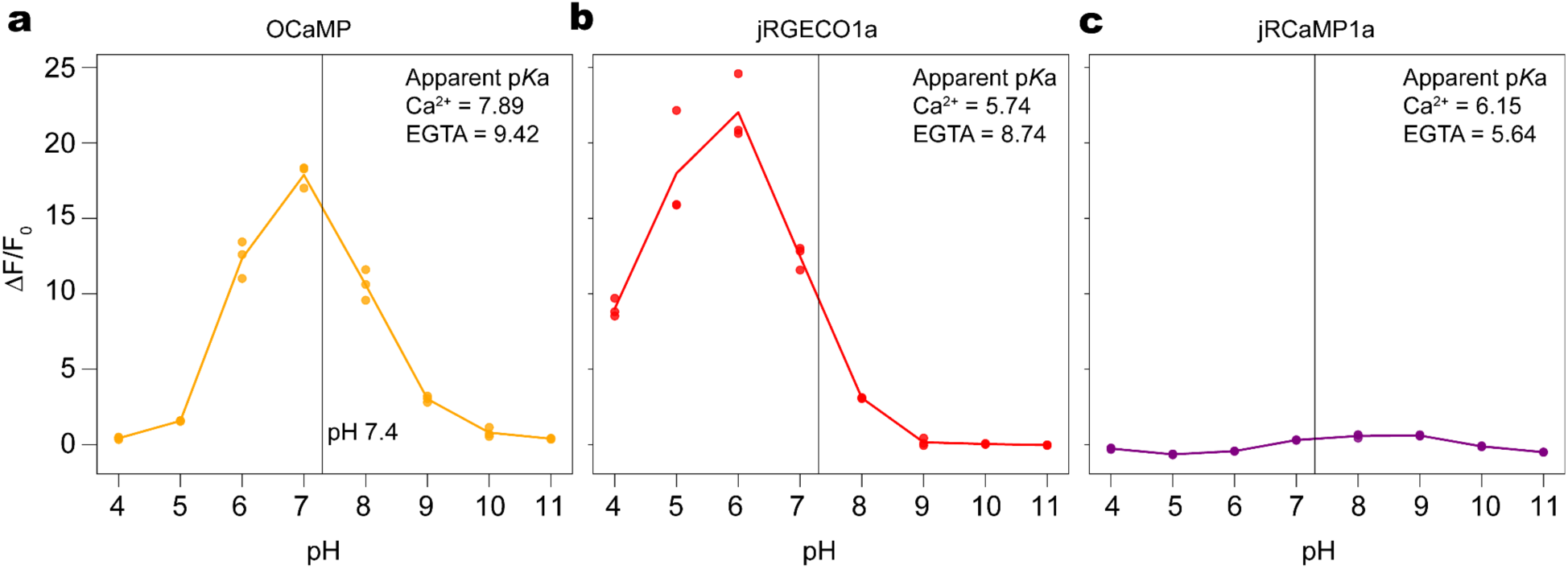
pH titration of purified soluble proteins. ΔF/F₀ responses as a function of pH for (a) OCaMP, (b) jRGECO1a, and (c) jRCaMP1a. Measurements were performed in Ca²⁺-bound conditions (10 mM Ca-EGTA buffer yielding 38 μM free Ca²⁺, buffered with 30 mM MOPS and 100 mM KCl) and Ca²⁺-free conditions (10 mM EGTA). Solid lines show mean values across three replicates (N = 3 titration series from a single protein preparation), with colored spheres representing individual data points. Apparent pKₐ values for each condition were obtained by fitting a sigmoidal curve to the Ca²⁺-bound and Ca²⁺-free conditions separately. A vertical black line at pH 7.4 marks physiological pH.

**Supplementary Figure 4.**
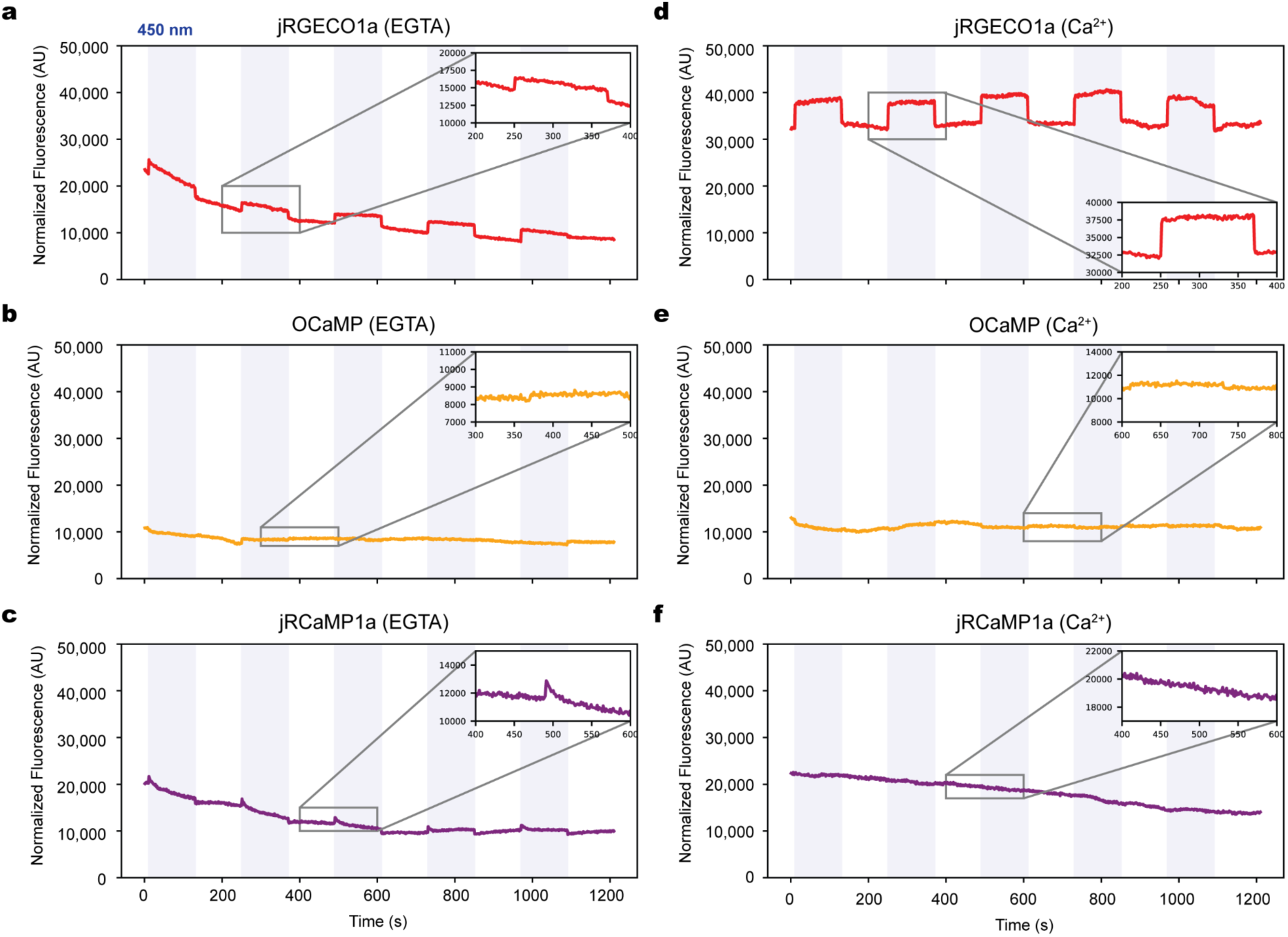
Photoactivation of jRGECO1a, OCaMP, and jRCaMP1a in EGTA and Ca²⁺ conditions. (**a–c**) HEK293 cells expressing jRGECO1a (**a**), OCaMP (**b**), or jRCaMP1a (**c**) were imaged in HHBSS buffer containing 5 mM EGTA. Cells were exposed to five 450 nm light pulses (0.0437 mW, 120 s duration) at regular intervals (gray shaded bars). Total imaging duration was 22 minutes. (**d–f**), Same experimental conditions as in **a–c**, but in the presence of 5 mM CaCl₂.

**Supplementary Figure 5.**
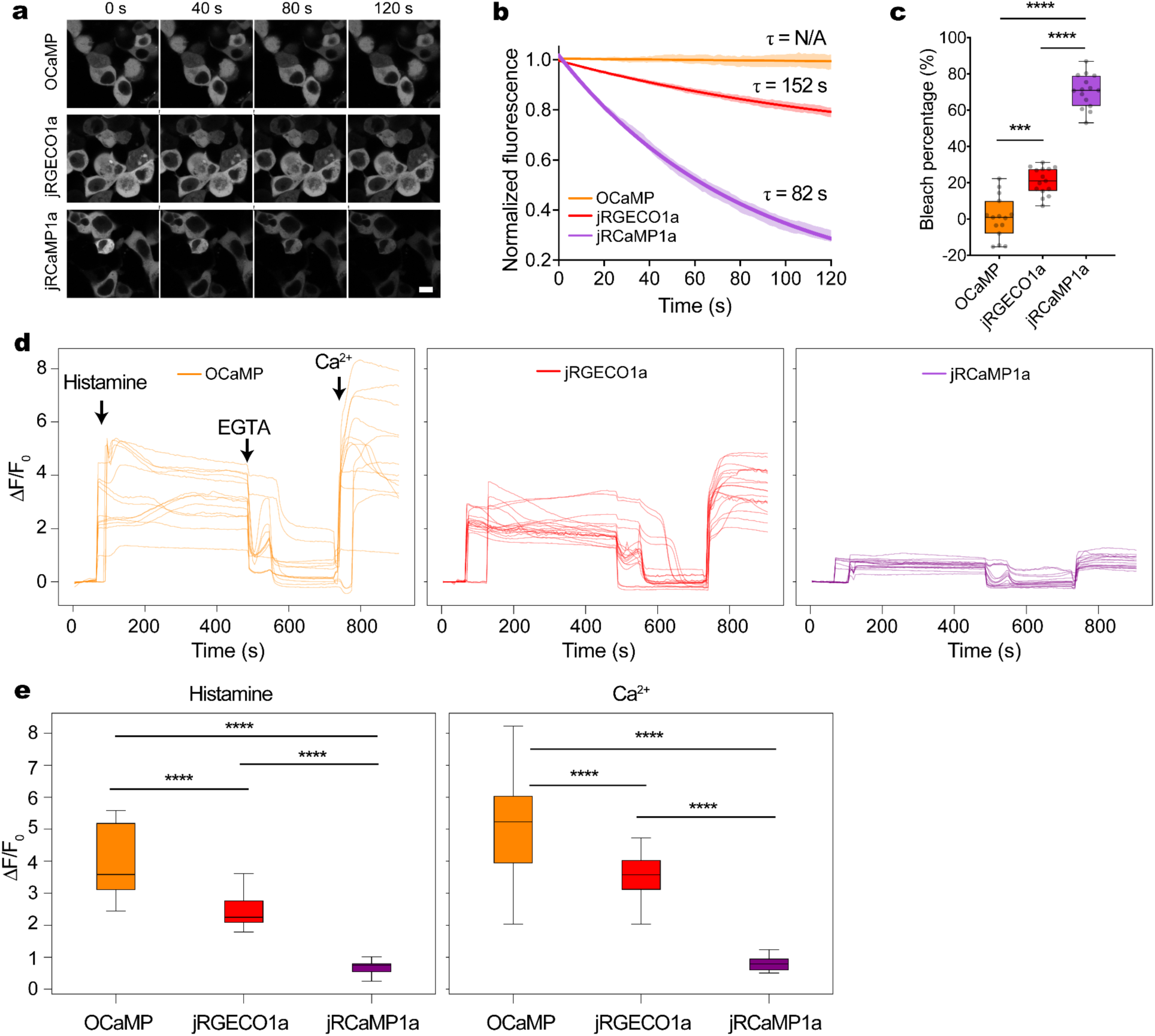
Photobleaching and ΔF/F₀ responses in mammalian cells. (a) Timelapse snapshots of HEK293 cells expressing OCaMP, jRGECO1a, and jRCaMP1a during continuous 1030 nm two-photon excitation. Scale bar: 10 μm. (b) Normalized fluorescence over time in HEK293 cells under 1030 nm excitation (n = 15 cells per sensor), baseline-normalized to t = 0. Solid line represents non-linear exponential decay fit; shaded area indicates ±SEM. (c) Bleach percentage in HEK293 cells, calculated as the average fluorescence at t = 120 s relative to fluorescence at t = 0. Tukey’s HSD: all pairwise comparisons p < 0.0001. (d) Individual ΔF/F₀ traces in HeLa cells following histamine stimulation under 545 nm one-photon excitation. (e) Quantification of peak ΔF/F₀ responses in HeLa cells under 545 nm excitation after histamine or calcium addition. For histamine stimulation: OCaMP (n = 20, median = 3.59, IQR = [3.12, 5.18]), jRGECO1a (n = 18, median = 2.25, IQR = [2.09, 2.77]), jRCaMP1a (n = 15, median = 0.75, IQR = [0.54, 0.79]). One-way ANOVA *p* = 1.48×10⁻¹⁷; Tukey’s HSD: all pairwise comparisons *p* < 0.0001 (). For calcium stimulation: OCaMP (n = 20, median = 5.23, IQR = [3.95, 6.03]), jRGECO1a (n = 18, median = 3.58, IQR = [3.12, 4.03]), jRCaMP1a (n = 15, median = 0.79, IQR = [0.60, 0.95]). One-way ANOVA *p* = 1.95×10⁻¹⁴; Tukey’s HSD: all pairwise comparisons *p* < 0.0001 ().

**Supplementary Figure 6.**
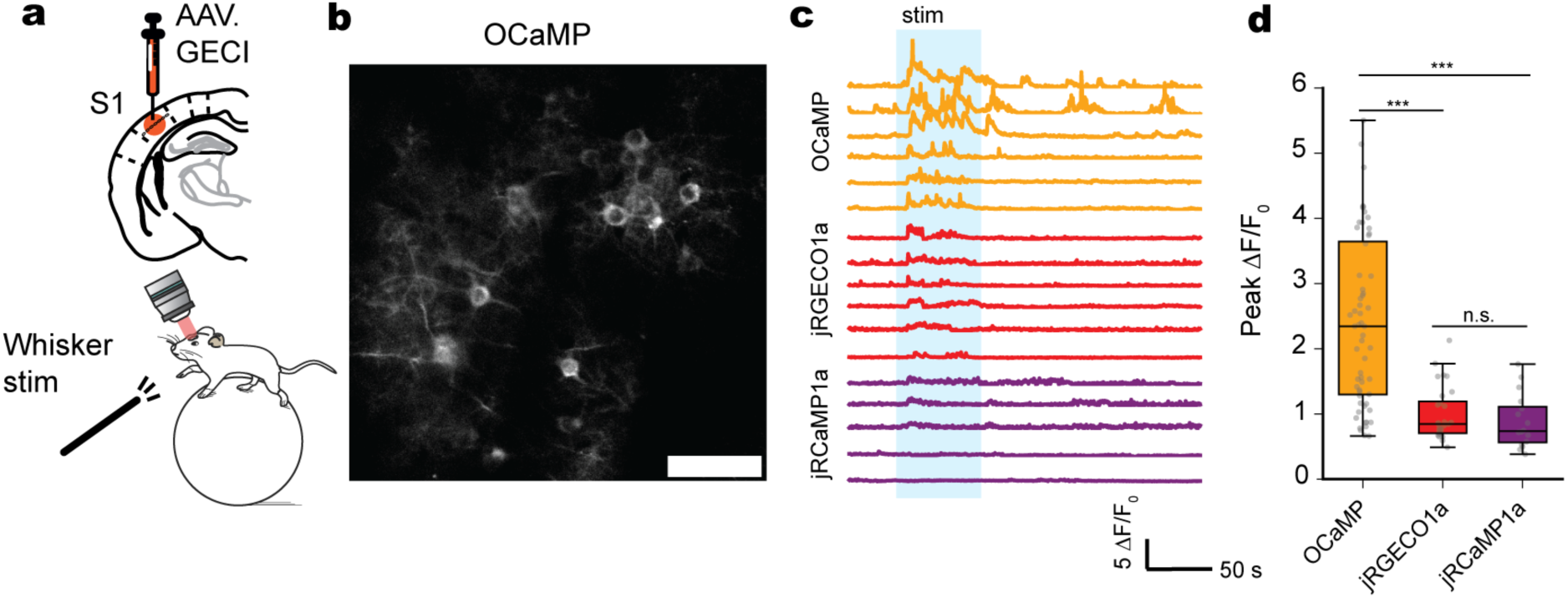
*In vivo* imaging of whisker-evoked calcium activity in mouse layer 2/3 barrel cortex neurons. (**a**) Experimental setup for awake two-photon imaging during contralateral whisker stimulation (left) and (**b**) representative single-plane average projection images from mice expressing OCaMP (right). Scale bar: 50 μm (OCaMP and jRGECO1a); 25 μm (jRCaMP1a). (**c**) Representative ΔF/F₀ traces from individual ROIs showing time-locked responses to whisker stimulation. (**d**) Peak ΔF/F₀ responses across individual ROI; OCaMP (*n* = 60 peaks), jRGECO1a (*n* = 28 peaks), and jRCaMP1a (*n* = 16 peaks). Data are shown as box and whisker plots (min to max values). Statistical significance was assessed using Brown-Forsythe ANOVA with Dunnett’s multiple comparisons test (n.s. = non-significant; \*\*\**P* < 0.0001).

## Methods

### Animal care and use statement

All experimental procedures involving animals were performed in accordance with protocols approved by the Institutional Animal Care and Use Committee at the respective institute (University of Calgary: AC22-0137, University of Geneva: GE253, GE358, GE388). This research has complied with all applicable ethical regulations.

C57BL/6J male mice (Charles River, Janvier Labs) were group housed on a 12-h light cycle with littermates under standardized conditions at the animal facilities at the University of Calgary or the University of Geneva until surgery.

All procedures were conducted in accordance with the guidelines of the Federal Food Safety and Veterinary Office of Switzerland and in agreement with the veterinary office of the Canton of Geneva (licence numbers GE253, GE358, GE388).

### Reagent and availability

Reagents are available by e-mail request or from Addgene (#s 224932, 224934, 224936, 225492, 229853)

### Molecular biology

Mammalian expression constructs encoding calcium indicators were synthesized (VectorBuilder) or generated by PCR amplification from existing templates. PCR products and destination vectors (pAAV.hSyn, pAAV.CAG, pAAV.hSyn.FLEX, or pAAV.FRT.FLEX) were digested with appropriate restriction enzymes or linearized by PCR and assembled using Gibson assembly (NEBuilder HiFi DNA Assembly Master Mix, New England Biolabs) following the manufacturer’s protocol. All constructs included an N-terminal RSET sequence derived from pRSET to enhance expression and solubility. Final plasmids were sequence-verified by Sanger sequencing.

### Construction of mammalian expression vectors

The mammalian expression vectors (Cre-independent pAAV.hSyn and pAAV.CAG, Cre-dependent pAAV.hSyn.FLEX, and FLP-dependent pAAV.FRT.FLEX) expressing the different Ca^2+^ indicators were commercially synthesized (VectorBuilder). The N-terminus contains the RSET peptide reader sequence carried over from pRSET for all sensors.

### AAV production and purification

AAV viral vectors containing the PHP.eB capsid were generated using the methods as previously described^28^. PHP.eB capsid has been engineered to efficiently transduce the central nervous system^28,29^. Briefly, 293FT cells (Thermo Fisher Scientific) were grown to about 90% confluence in hyperflasks (Corning) and cotransfected with 129 μg pHELPER (Agilent), 238 μg rep-cap plasmid encoding PHP.eB pUCmini-iCAP-PHP.eB was a gift from Viviana Gradinaru (Addgene plasmid # 103005, RRID:Addgene_103005)), and 64.6 μg of transfer plasmid using the PEIpro transfection reagent (Polyplus). AAVs were precipitated from medium harvested after 3 days and 5 days using 40% PEG/2.5 M NaCl in buffer containing 500 mM NaCl, 40 mM Tris base, and 10 mM MgCl_2_. The lysate was incubated with 100 U/mL salt-active nuclease (Arcticzymes) at 37°C for 1 hour and then centrifuged at 2,000*g* for 15 minutes. AAV was purified from the resulting lysate using an iodixanol step gradient containing 15%, 25%, 40%, and 60% iodixanol in OptiSeal tubes (Beckman Coulter) followed by centrifugation at 350,000*g* using a Type 70 Ti ultracentrifuge rotor (Beckman Coulter). After centrifugation, the AAVs were harvested from the 40% layer using a 10 mL syringe and 16-gauge needle, diluted in 1× PBS containing 0.001% Pluronic F68 (Gibco), and filtered using a 0.2 μm syringe filter. The AAVs were concentrated and buffer-exchanged by 5 rounds of centrifugation using Amicon Ultra-15 100-kDa molecular weight cutoff centrifugal filter units (MilliporeSigma). The titer was determined using the qPCR Adeno-Associated Virus Titration kit (Applied Biological Materials), and the purity was verified by SDS-PAGE and total protein staining using ReadyBlue instant gel stain (Sigma).

### Protein purification and *in vitro* characterization

Constructs were subcloned into a pRSET vector containing an N-terminal His tag and RSET peptide. T7Express (NEB) *E. coli* were transformed and cultured in LB media at 37°C until OD₆₀₀ reached 0.6–0.8. Cells were lysed by sonication in lysis buffer (50 mM Tris-HCl pH 7.5, 300 mM NaCl, 10 mM imidazole, 1 mM PMSF), and lysates were cleared by centrifugation (20,000 × g, 30 min). Proteins were purified by Ni²⁺-NTA affinity chromatography (Qiagen), washed with buffer containing 10 mM imidazole, and eluted with 250 mM imidazole. Eluted fractions were concentrated and buffer-exchanged into TBS (50 mM Tris-HCl, 100 mM NaCl, pH 7.2) using a 10 kDa MWCO centrifugal filter (Amicon). Protein concentration was determined by absorbance at 280 nm (Nanodrop, Thermo Scientific).

### One-photon photophysical measurements

One-photon fluorescence intensity spectra in saturated calcium (10 mM [Ca²⁺] buffered in TBS) and calcium-free (TBS only) conditions were obtained using a Tecan Infinite M1000 plate reader. For OCaMP and OCaMP-fast, excitation scans were collected from 450 to 580 nm (emission set at 600 nm, bandwidth 10 nm, step 2 nm), and emission scans were collected from 500 to 700 nm (excitation set at 530 nm, bandwidth 10 nm, step 2 nm). For jRGECO1a, jRCaMP1a, and XCaMP-R, excitation scans were collected from 450 to 580 nm (emission set at 600 nm, bandwidth 10 nm, step 2 nm), and emission scans were collected from 500 to 700 nm (excitation set at 560 nm, bandwidth 10 nm, step 2 nm). For jYCaMP1a, excitation scans were collected from 420 to 550 nm (emission set at 530 nm, bandwidth 10 nm, step 2 nm), and emission scans were collected from 460 to 650 nm (excitation set at 500 nm, bandwidth 10 nm, step 2 nm).

Buffers for calcium affinity titration, with free [Ca²⁺] ranging from 0 to 39 μM, were prepared using the Calcium Calibration Buffer Kit #2 (ThermoFisher) and mixed 1:1 with diluted protein samples. Single-point excitation intensities were recorded by exciting at 545 nm and collecting emission at 565 nm for OCaMP and OCaMP-fast, 570 nm excitation and 600 nm emission for jRGECO1a, jRCaMP1a, and XCaMP-R, and 530 nm excitation and 550 nm emission for jYCaMP1a (all with 10 nm bandwidth).

Extinction coefficients were determined using the alkali denaturation method. Proteins were denatured in 0.1 M NaOH and absorbance of the denatured chromophore was measured at 446 nm, using the extinction coefficient of denatured fluorescein as a reference (ε = 44,000 M⁻¹ cm⁻¹ at 446 nm). Extinction coefficients for the native state were calculated by dividing the native absorbance by the concentration determined from the denatured state. Quantum yield (QY) measurements were performed using a Quantaurus-QY spectrophotometer (Hamamatsu Photonics, Hamamatsu City, Japan) for the various indicators in TBS and in 10 mM [Ca²⁺].

For pH titrations, purified proteins were diluted into a series of MOPS-buffered solutions (30 mM MOPS, 100 mM KCl) spanning pH 4 to 11, either in 10 mM EGTA (Ca²⁺-free) or in 10 mM Ca-EGTA buffer yielding 38 μM free Ca²⁺ (Ca²⁺-bound). Fluorescence was recorded at fixed excitation and emission wavelengths: 545/566 nm for OCaMP and 570/600 nm for jRGECO1a and jRCaMP1a (excitation/emission). ΔF/F₀ was calculated from the fluorescence values, and apparent pKₐ values were obtained by fitting a sigmoidal curve to each condition.

### Stopped-flow kinetics

Measurements of OFF kinetics were made using an applied Photophysics SX20 Stopped-flow Reaction Analyzer using fluorescence detection, exciting at 530 nm and detecting with a 550 nm long-pass filter in a 10 mm quartz flow cell at 22 °C. The dead time of the instrument was 1.1 ms. Specifically, 2 µM of each calcium indicator in 1 mM [Ca^2+^] (TBS solution, pH 7.2) was rapidly mixed at 1:1 ratio with 10 mM of EFTA (TBS, pH 7.2) at room temperature. At least three to five replicates were collected per indicator, averaged, and a monoexponential decay curve was fit to the average trace and k_off_ values were determined (units of s^-1^) using Prism GraphPad.

### Two-photon photophysical measurements

Two-photon measurements were made using an inverted microscope (IX81, Olympus), equipped with a 60X, 1.2 NA water immersion objective (Olympus). The various calcium indicators were measured in 30 mM MOPS buffers either with 39uM free Calcium (+Ca) or 0uM free Calcium (-Ca) buffered at pH 7.2. Excitation was performed with an 80 Mhz Ti-Sapphire laser (Chameleon Ultra II, Coherent) to collect 2P excitation spectra from 710 to 1,080 nm and with an OPO(Chameleon Compact OPO, Coherent) for 1000-1250nm. Fluorescence collected by the objective was passed through a shortpass filter (720SP, Semrock) and a band pass filter (539BP278, Semrock), and detected by a filter-coupled Avalanche Photodiode (APD) (SPCM_AQRH-14, Perkin Elmer). The brightness spectra were normalized to 1 µM concentration and further used to obtain action cross-section spectra (AXS) with Rhodamine B dye^30^ as a reference.

### Preparation of HEK239 cells

HEK293 cells were grown in Dulbecco’s Modified Eagle’s Medium - high glucose (Sigma-Aldrich) (DMEM) with 10% FBS (Sigma-Aldrich) and penicillin streptomycin at 37°C. Cells were plated on poly-D-lysine coated, glass bottom (#1.5 coverslip) 24-well plate, and transfected with pAAV.CAG.OcaMP, pGP-CMV-NES-jRGECO1a and pGP-CMV-NES-jRCaMP1a, 1.5 µg DNA per well, in three replicates, using the Lipofectamine™ 3000 Transfection kit (Invitrogen) standard protocol. Cells were grown at 37°C for 48 h before imaging.

### Characterization and analysis in HeLa cells

The cells were cultured in DMEM medium (Nacalai Tesque, 08456-65) supplemented with 10% FBS (Sigma-Aldrich, F7524-500ML) and 1% penicillin–streptomycin (Nacalai Tesque, 09367-34). HeLa cells (ATCC CCL-2, ∼ 5 x 10^4^/dish) were seeded into 35-mm glass-bottom cell-culture dishes (Iwaki, 3911-035) with 200 μl Opti-MEMI (Thermo Fisher Scientific, 31985-070) and transfected with 2 μg of plasmid using 4 μl polyethyleneimine (Polysciences, 24765-1, 1 mg polyethyleneimine was diluted in 1 ml ultrapure water). Transfected cells were imaged 48-72 hours after transfection using a IX83 wide-field fluorescence microscopy (Olympus) equipped with a pE-300 LED light source (CoolLED), a 40× objective lens (numerical aperture = 1.3; oil), an ImagEM X2 EM-CCD camera (Hamamatsu photonics), Cellsens software (Olympus) and a STR stage incubator (Tokai Hit). The filter sets for imaging were excitation 545/20nm, dichroic mirror 565-nm dclp, and emission 598/55 nm, respectively. For imaging of Ca^2+^-dependent fluorescence, the dish (Iwaki, 10-mm glass-bottom) was washed twice with Hank’s balanced salt solution (+) (HBSS(+); Nacalai Tesque, 09735-75), and the buffer was exchanged to 1 ml HBSS(+) supplemented with 10 mM HEPES (Nacalai Tesque, 17557-94)) at 37 °C just before the imaging. Then histamine (Wako, 087-03533, 50 μM final concentration) was added one minute after imaging started. At the indicated time, imaging was stopped, and the dish was washed twice with HBSS without Ca^2+^ (HBSS(-)) (Nacalai Tesque, 17461-05), and 1 mL of HBSS(-) was added. Then, EGTA/ionomycin (2.5 mM/1.5 μM final concentration) were added to quench the Ca^2+^-dependent oscillation. At the indicated time, imaging was stopped again, and the dish was washed twice more and supplemented with HBSS(-). After resuming, Ca^2+^/ionomycin (5 mM/1.5 μM final concentration) was immediately added to saturate the indicator.

### Two-photon photobleaching in HEK293 cells

Transfected HEK293 cells in HHBSS (HBSS media (Gibco) supplemented with HEPES to 10 mM) were exposed to continuous, two-photon, 1030 nm light (20 mW/cm^2^) for 120 s. Imaging was done at 40x magnification with a frame rate of 0.775 Hz. Images were analyzed using Fiji, a nonlinear decay curve and tau values were fitted and obtained using Prism Graphpad.

### Photoswitching in HEK293 cells

Transfected HEK293 cells in HHBSS were imaged continuously at 570 nm (for jRGECO1a and jRCaMP1a) and 545 nm (for OCaMP) for 22 minutes with five 2 minute long pulses of 450 nm light (0.0437 mW) starting at minute 2 and then every 4 minutes. Imaging was done under a 10x objective and zoom factor 3, with a frame rate of 0.775 Hz. To chelate Ca^2+^, EGTA was added together with 1.5 µM ionomycin. For saturating the sensors, CaCl₂ was added to 5 mM together with 1.5 µM ionomycin. Fluorescence values were obtained using FIJI.

### Preparation of neuronal cultures

Neonatal rat pups (P0) were euthanized; cortical and hippocampal tissue was dissected, and dissociated in papain enzyme (Worthington Biochemicals, ∼35 U/cortical and hippocampal pair) in neural dissection solution (10 mM HEPES pH 7.4 in Hanks’ Balance Salt Solution) for 30 minutes at 37 °C. After 30 minutes, enzyme solution was aspirated out and tissue pieces were subjected to trituration in 10% fetal bovine serum containing MEM media. Following trituration, cell suspension was filtered through a 40 μm strainer, and resulting single cell suspension was centrifuged. Cell pellet was resuspended in plating media (28mM glucose, 2.4mM NaHCO3, 100µg/ml transferrin, 25µg/ml insulin, 2mM L-glutamine, 10% fetal bovine serum in MEM) and cell counts were taken. Electroporation was conducted using the Lonza/Amaxa 4D nucleofector according to the manufacturer’s instructions, using 500 ng of plasmid and 5×105 viable cells per transfection. Transfected cells were then seeded into 3 replicate wells in poly-D-lysine-coated glass-bottom 24-well plates or 35 mm dishes, and cultured at 37 °C with 5 % CO2. Cultures were fed twice a week by replacing 50% of the medium with fresh NbActiv.

### Neuronal stimulation

Calcium indicators were cloned into a mammalian expression vector (pAAV with hSyn promoter) and transfected into hippocampal and/or cortical primary cultures from neonatal (P0) pups. Imaging was performed in poly-D-lysine-coated 35 mm glass-bottom dishes. 12-14 days post transfection, culture medium was exchanged with 1 mL imaging buffer (145 mM NaCl, 2.5 mM KCl, 10 mM glucose, 10 mM HEPES, 2 mM CaCl_2_, 1 mM MgCl_2_, pH 7.3) and imaged in 1 mL imaging buffer and drug cocktail to inhibit synaptic transmission (10 μM CNQX, 10 μM (R)-CPP, 10 μM gabazine, 1 mM (S)-MCPG (Tocris)). Neurons were field stimulated with 1, 3, 5, 10, 20, 40, 80, and 160 pulses at 33.3 Hz (30 ms exposure time) and imaged with 10X objective. The indicators were excited at 555 nm for OCaMP, OCaMP-fast, jRGECO1a, jRCaMP1a, and XCaMP-R (emission 605/50 nm using RFP filter cube), and at 505 nm for jYCaMP1 (emission 535/30 using YFP filter cube). Imaging was performed at room temperature. Analysis of recordings was performed with custom Python code.

### Characterization in primary rat neuronal cultures

Regions of interest (ROIs) were manually drawn in FIJI and fluorescence traces were extracted. Baseline fluorescence (F₀) for each trace was computed as the mean of the 10 frames preceding stimulation. ΔF/F₀ traces were calculated for each ROI and peak amplitudes were quantified per stimulus condition. Mean and standard error of the mean (SEM) were calculated across ROIs for each indicator and plotted using custom R scripts.

ON and OFF kinetics were extracted from ΔF/F₀ traces using custom Python code. For ON kinetics, rise times (10–90%) were calculated by identifying the frame of stimulus onset and measuring the time required to reach 90% of the peak ΔF/F₀. For OFF kinetics, single-exponential decay curves were fit to the falling phase of the ΔF/F₀ signal following the stimulus, and decay time constants (τ) were obtained using least-squares fitting in Python.

### Expression and Functional Imaging Analysis in Zebrafish Larvae

Transgenic expression of genetically encoded calcium indicators was achieved by co-injecting Tol2 transposase mRNA (25 ng/μL) with a Tol2-based Elavl3-driven expression construct (40 ng/μL) into one-cell stage zebrafish embryos. Injected embryos were raised in E3 medium at 28.5 °C and imaged at 5 days post-fertilization (dpf). For imaging, larvae were embedded dorsal side up in 1.5% low-melting point agarose and maintained in standard E3 solution.

Spontaneous neuronal activity was recorded using one-photon (1P, 545 nm excitation) and two-photon (2P, 1030 nm excitation) fluorescence microscopy at a frame rate of 3 Hz. Image processing and fluorescence analysis were performed using custom Python scripts. Registered image stacks were first mean-projected to assist in ROI detection. ROIs were identified using skimage.feature.peak_local_max (scikit-image v0.22.0) on Gaussian-smoothed images (σ = 1.0, minimum distance = 6 pixels). Circular ROIs (radius = 5 pixels) were used to extract raw fluorescence traces. ΔF/F₀ was computed using a rolling baseline defined as the 10th percentile over a 10-frame window. ROIs with maximum ΔF/F₀ below 0.1 were excluded. The top 10 ROIs from each file were selected based on peak ΔF/F₀ amplitude. Distributions of ΔF/F₀ peaks were summarized using box plots. Analyses were performed using NumPy (v1.26.4), SciPy (v1.13.0), and scikit-image (v0.22.0), and plots were generated using Matplotlib (v3.8.4).

### Surgical procedures and viral injections (University of Genève)

Male C57BL/6 mice 8-13 weeks old were anesthetized with a mix of O_2_ and 4% isoflurane at 1-2 L min^−1^ to induce anaesthesia followed by an intraperitoneal injection of MMF solution, consisting of 0.2 mg kg^−1^ medetomidine (Dormitor, Orion Pharma), 5 mg kg^−1^ midazolam (Dormicum, Roche), and 0.05 mg kg^−1^ fentanyl (Fentanyl, Sinetica) diluted in sterile 0.9% NaCl.

Toe pinch reflexes were tested 15 min after anaesthetic injection to ensure anaesthesia depth. Dexamethasone (Mephameson, mepha) 0.02 ml at 4 mg/ml was administered by intramuscular injection to the quadriceps which reduced cortical stress response during the surgery.

The head of the mouse was shaved, then the mouse was placed on a heating blanket and the head stabilized in a stereotaxic apparatus. The eyes were protected from dehydration and irritation by application of an eye-ointment. The scalp was washed with betadine, then lidocaine (Streuli) 1% was injected under the skin before scissors were used to remove a flap of skin of approximately 1 cm squared covering the skull. The fascia was gently scraped from the skull with a swab. The area of interest was marked with a pen. The surface of the skull was drilled to increase the surface of adhesion with dental cement. A circular area (diameter 3 mm) of the skull overlying one hemisphere was removed using a 3-mm biopsy puncher exposing the underlying brain. Sterile cortex buffer (125 mM NaCL, 5 mM KCL, 10 mM glucose, 1M CaCl_2_, 1M MgSO_4_) was applied to the surface of the brain. Dexamethasone (0.02 ml at 4 mg/ml) was applied topically on the surface of the craniotomy. Viruses were delivered to L2/3 of the right barrel cortex in S1 at the approximate location of the C2 barrel-related column (1.4 mm posterior, 3.5 mm lateral from bregma, 300 µm below the pia). Viruses were injected mechanically (a volume of 100-200 nl), slowly over the time course of 1 to 2 min using an oil hydraulic manipulator system (MMO-220A, Narishige) with a 20-μm diameter glass capillary.

Viral injections were performed using genetically encoded indicators delivered through Adeno-Associated Viruses (AAV). For OCaMP experiments, either PHPEB.hSyn.OCaMP (∼5×10¹² GC/mL) was injected either at full titer, diluted 2x, 10x, 50x 100x or AAV1.hSyn.FLEX.OCaMP (∼5×10¹² GC/mL) was injected at full titer mixed with AAV1.mCaMKIIα.iCre.WPRE-hGHp(A) (VVF/ETHZ # v206-1, 8.7*10^12^ GC/mL) diluted 10x or 100x. For jRGECO1a experiments, AAV1.syn.FLEX.NES-jRGECO1a.WPRE.SV40 (Addgene plasmid # 100852, 2.2×10^13^ GC/mL) was injected at full titer mixed with AAV1.mCaMKIIα.iCre.WPRE.hGHp(A) (VVF/ETHZ # v206-1, 8.7*10^12^ GC/mL) diluted 50x or 100x.

For jRCaMP1a experiments, AAV1.Syn.NES.jRCaMP1a.WPRE.SV40 (Addgene plasmid #100848, ≥ 1×10¹³ GC/mL) was diluted 10x or 100x.

The pipette was left in place for 3–5 min before removing it from the brain. A coverslip of ∼ 100 μm thickness and a 3 mm diameter was placed onto the brain. The optical window was sealed to the skull with cyanoacrylate adhesive (Loctite 406 or Pattex Ultra Gel) and dental acrylic (two phase component: Jet Denture Repair Powder and Jet Liquid) covering all exposed areas of the skull and edges of the cover glass. All wound margins were sealed with dental acrylic. A custom-built stainless steel 1.6 g or aluminum bar 0.54 g was embedded into the dental acrylic over the intact hemisphere. The anaesthetic mix (AFB) was antagonized by a waking solution injected subcutaneously: atipamezole (Antisedan Atipamezol) 2.5 mg/kg, flumazenil (Labatec-pharma) 0.5 mg/kg, buprenorphine (Bupaq Buprenorphium) 0.1 mg/kg. During the waking process mice were placed individually in a recovery cage for approximately one hour on a heating pad and given moistened food. Mice were given the analgesic carprofen (Rimadyl, Zoetis) 5 mg/kg twice per day during the 48 hours following the surgery and monitored daily to ensure full recovery.

In addition, a second anti-inflammatory Ibuprofen (Algifor, Verfora) 10 ml/100 ml was added to drinking water for 3-days post-surgery. Animals were then put back in their home cage to recover from the surgery.

### 2-photon laser scanning microscopy (University of Genève)

Mice were handled and accustomed to be head restrained on the training setup for 10–15 min 2-3 days after a minimum period of 15 days post-surgery and viral transfection. The acquisitions used for the systematic comparison of the Ca^2+^ indicators were acquired between 20 and 45 days after AAV injection with most acquisitions done between 26- and 32-days post-injection.

We used a custom built 2-photon laser scanning microscope mounted onto a modular *in vivo* multiphoton microscopy system (https://www.janelia.org/open-science/mimms-10-2016) equipped with an 8-kHz resonant scanner and a 25x 1.1 NA objective (Nikon, N25X-APO-MP). Fluorophores were excited using a Ti:Sapphire laser (Chameleon Ultra, Coherent) tuned to λ = 1020 nm. Fluorescent signals were collected through the detection path consisting of an emission filter ET620/60 m (Chroma), a dichroic mirror (565dcxr, Chroma) and a GaAsP photomultiplier tubes (10770PB-40, Hamamatsu). Images were acquired with Scanimage 2016b^31^. Functional images of L2/3 cells (512 × 512 pixels, 225 x 225 μm) and L1 neuropil, 50– 100 μm under the pia mater (256 × 128 pixels,130 x 65 μm) were collected at 30 Hz or 60 Hz, respectively with a laser power between 41 and 55 mW measured at the front aperture of the objective. All acquisitions lasted 2 minutes. Photobleaching acquisitions of L1 neuropil, 50-100 μm under the pia mater (256 × 128 pixels, 67 x 33.5 μm) were collected at 60 Hz with a laser power between 59 and 66 mW.

### Image Processing (University of Genève)

Time-series were motion-corrected and segmented with the Suite2p toolbox32. Detected ROIs were manually selected or discarded based on their shape. Somatic ROIs were kept if they were donut-shaped or showed clear activity, while dendritic and axonal ROIs were included if they exhibited dendrite- or axon-like morphology or demonstrated clear activity. The fluorescence time course of L2/3 cell bodies was measured by averaging all pixels within individual ROIs, and all subsequent analyses were performed using custom MATLAB scripts (The MathWorks, Natick, MA, USA). For OCaMP and jRGECO1a acquisitions, but not for dendrites and axons of L1, neuropil signal was subtracted from the raw fluorescence of L2/3 cell bodies as F_neuron = F_raw – 0.7 × F_neuropil, while jRCaMP1a, which was more sparsely expressed, was not neuropil-corrected.

Normalized calcium traces (ΔF/F₀) were calculated as (F−F₀)/F₀, where F₀ was defined as the 30th percentile of the individual mean baseline fluorescence over a 10-second rolling window. The fluorescence signals F(t) were then converted to z-scores for peak extraction of spontaneous activity, and traces were additionally filtered using a Savitzky-Golay function (2nd order, 366-ms span). Local maxima were identified in the z-scored traces for each ROI using the findpeaks function, with an adaptive threshold set as median(X) + 2 × MAD(X) + STD(X), where X represents the Ca²⁺ trace in z-scores and MAD is calculated as mean(abs(X−mean(X))). The minimum peak distance was set to 300 ms, minimum peak prominence was set to the adaptive threshold, and peaks were required to be at least 5 data points apart to discard artifacts.

Cell bodies of ROIs identified by Suite2p with at least two events during the 2-minute recording were classified as responsive. The total number of neurons expressing the indicator in each field-of-view was counted in a semi-automated manner: most cell bodies were detected using the anatomical detection of Suite2p, and any visually identified missing cells were manually added. To measure the decay time constant (τ), a monoexponential function, f(t) = A·e×(−bt), where A is amplitude, b is decay rate, and t is time, was fitted to each detected peak. The decay time constant τ was then calculated as 1/b, and only fits with an R-squared value above 0.7 were retained for measurements.

### Surgical procedures and viral injections (University of Calgary)

All procedures for awake two-photon imaging were performed according to previously published protocols^33^. Briefly, mice were anesthetized with isoflurane and received subcutaneous injections of buprenorphine (0.1 mg/kg), enrofloxacin (10 mg/kg), and meloxicam (3 mg/kg). Under aseptic conditions, the scalp was removed, the skull was cleaned, and a U-shaped metal head bar was affixed to the skull using fast glue and dental cement (Tetric EvoFlow, Patterson Dental, Calgary, AB). A craniotomy was performed over the barrel cortex under continuous immersion in HEPES-buffered artificial cerebrospinal fluid (142 mM NaCl, 5 mM KCl, 10 mM glucose, 10 mM HEPES, 3.1 mM CaCl₂, 1.3 mM MgCl₂). AAV vectors encoding calcium indicators were injected into the barrel cortex. After injection, the craniotomy was sealed with a t-shaped glass coverslip.

### Mouse Training (University of Calgary)

Four weeks after surgery, mice were habituated to head-fixation while running on an air-supported spherical Styrofoam ball. Mice were initially allowed to run freely for 15 minutes, after which intermittent air-puff stimulation was delivered to the contralateral whiskers (15 stimulations, 5 seconds each, every 60 seconds).

### Two-photon laser scanning microscopy (University of Calgary)

Imaging was performed using a custom-built two-photon microscope powered by a Ti:Sapphire laser (Coherent Ultra II, ∼4 W average output at 800 nm, ∼80 MHz repetition rate). Image acquisition was controlled using ScanImage software (version 3.81, HHMI/Janelia Research Campus). Excitation was performed at 1030 nm for OCaMP and 1040 nm for jRGECO1a and jRCaMP1a. The objective used was a 16x water-immersion, 0.8 NA (Nikon). The microscope was equipped with a primary dichroic mirror at 695 nm. Emitted fluorescence was further split and filtered using either a 560 nm secondary dichroic mirror and a 605/70 nm bandpass filter (for jRGECO1a and jRCaMP1a) or a 550 nm secondary dichroic and a 585/65 nm bandpass filter (for OCaMP) (Chroma Technologies).

### Sensory stimulation and image analysis (University of Calgary)

Neuronal responses to contralateral whisker stimulation were recorded. After a 30-second baseline recording, air puffs were delivered to the contralateral whiskers through two glass tubes for 30 seconds. Neuronal calcium signals were analyzed using custom Matlab scripts^33,34^. Regions of interest (ROIs) corresponding to individual neuronal somata were selected, and ΔF/F₀ traces were calculated by normalizing the fluorescence signal (F) to the baseline fluorescence (F₀) defined as the average signal during a pre-stimulation period. Peak ΔF/F₀ responses were identified within the stimulation window using a prominence-based peak detection algorithm (SciPy’s find_peaks function, prominence threshold of 0.5). For each detected peak during stimulation, ΔF/F₀ values were extracted and pooled across ROIs for statistical analysis.

### Surgical procedures and viral injections (Simultaneous two-photon imaging and juxtacellular recordings in mouse barrel cortex)

All animal procedures were performed in accordance with the protocols approved by the ethics committee of University of Geneva and the authorities of Canton of Geneva (license number GE253, GE388). Male adult mice of C57BL/6J genetic background (Charles River) of 5 or 17 weeks were used. To express calcium indicators at the mouse barrel cortex, stereotaxic injection of viral vectors was performed when the mice were under anesthesia, induced by a mixture of oxygen and 4% isoflurane (Provet) at 0.8 L/min and maintained by intraperitoneal injection of MMF solution, consisting of 0.2 mg/kg medetomidine (Orion Pharma), 5 mg/ midazolam (Sintetica), and 0.05 mg/kg fentanyl (Mepha) diluted in sterile 0.9% NaCl.

Viral vectors of PHPEB.hSyn.OCaMP were used for expressing OCaMP and pAAV.Syn.Flex.NES-jRGECO1a.WPRE.SV40 (AAV1) (Addgene plasmid # 100853) in combination with ssAAV-1/2-mCaMKIIa-iCre-WPRE-hGHp(A) (VVF/ETHZ # v206-1) for expressing jRGECO1a. Mice were head-fixed on a stereotaxic apparatus (Stoelting). Mouse body temperature was maintained at ∼37 °C. Eye ointment was applied to prevent dehydration. Analgesia was provided by local application of 1% lidocaine (Streuli). Betadine and 70% ethanol were applied at the skin of mouse head for disinfection. After removing the skin at the skull, a craniotomy was made over the right barrel cortex (1.4 mm posterior to bregma, 3.5 mm lateral to midline) using the dental drill. ∼100-200 nL virus was infused in 1-2 sites at ∼300 µm deep (layer 2/3) at a speed of ∼60 nL/min. Cortex buffer (125 mM NaCl, 5 mM KCl, 10 mM glucose, 10 mM HEPES, 2 mM CaCl_2_, 2 mM MgSO_4_) or Ringer’s solution solution (145 mM NaCl, 5.4 mM KCl, 10 mM HEPES, 1 mM MgCl_2_, and 1.8 mM CaCl_2_) was used for keeping the brain hydrated or washing the wound. After wound closure using surgical glue and suture, mice were put back to home cage for recovery.

Acute experiments of simultaneous two-photon imaging and electrophysiology were carried out ∼3 weeks to 1.5 months after viral injection for OCaMP and ∼2-2.5 months after viral injection for jRGECO1a. Following the same surgical procedures described above until the skull was exposed, a metal plate (∼2*3 cm^2^) was attached at the right mouse barrel cortex using cyanoacrylate glue and dental cement. A circular craniotomy of ∼1-2 mm diameter was made and the dura mater was removed. Agarose (0.5-2% in Ringer’s solution) and a cover glass were applied on top of craniotomy to dampen the tissue movement.

Experiments were performed with a custom-built two-photon (2P) laser scanning microscope (Holtmaat et al., 2009, Nat Protoc), equipped with a Ti:Sapphire laser (Chameleon Ultra, Coherent), a Pockel cell (Conoptics), galvanometric scan mirrors (Cambridge Technology), GaAsP photomultiplier tubes (10770PB-40, Hamamatsu), and a 16x water-immersion objective (0.8 NA, CFI75, Nikon). A laser wavelength of 1020 nm at a power of ∼69-124 mW (measured at the front aperture of objective) was used for exciting OCaMP or jRGECO1a. Emitted light was spectrally separated via a 565 nm dichoric mirror (565dcxr, Chroma) and two band-pass filters emission filters (510/50 nm and 620/60 nm; Chroma). Imaging acquisition was controlled via ScanImage software (Pologruto et al., 2003, BioMedical Engineering OnLine). Functional images (64*64 pixels and 53*53 µm^2^) of layer 2/3 cells (∼79-194 µm under the pia mater) were acquired at a frame rate of ∼30 Hz.

Juxtacellular recordings of layer 2/3 neurons were obtained using 2P-guided patching (Kitamura et al., 2008, Nat Methods). Patch pipettes of 5-8 MΩ tip resistance were fabricated from borosilicate glass by using a vertical microelectrode puller (Narishige). Ringer’s solution was used for both internal solution in patch pipette and external solution at the craniotomy. AlexaFlour-488 (30 µM) was added in the internal solution for visualizing the pipette during 2P imaging. After obtaining a juxtacellular recording (initial seal resistance ranged 9-91 MΩ, n=26 cells), spontaneous spiking activity was measured at current-clamp or voltage-clamp configuration while simultaneous functional imaging. Electrophysiological data were filtered at 6-10 kHz, sampled at 20 kHz using a Multiclamp 700B amplifier (Axon Instruments) and a Digidata 1440A digitizer (Molecular Devices). Data acquisition of electrophysiological recordings were controlled by pCLAMP 10 software (Molecular Devices), which could send output to ScanImage software for triggering the acquisition of functional imaging.

### Data analysis

2P images were processed using ImageJ (https://imagej.net/ij/) and custom-written scripts in MATLAB (MathWorks). Lateral drifts of 2P images were corrected based on the OCaMP (/jRGECO1a) signal using the NoRMCorre toolbox in MATLAB. Regions of interests (ROIs) covering the patched cell soma were manually selected based on calcium signal using ImageJ. The fluorescence signal of each ROI was obtained by averaging all pixels within the ROI, while neuropil signal around the soma was not subtracted.

Based on electrophysiological recordings, AP events were identified as 1) spikes of inter-spike interval<0.2 s (verified also by visual inspection); 2) inter-event interval (according to the first AP of neighboring AP events)>1 s.

The time-series of calcium signal was normalized as ΔF/F_0_=(F(t)-F_0_)/F_0_. For calcium signal of an AP event, F_0_ was calculated as the fluorescence average across the time from -11 to -1 image frame to the first AP peak; for a longer time-series of calcium signal including multiple AP events, F_0_ was calculated as the 10^th^ percentile. To determine the amplitudes and kinetics of calcium signal vs AP number, peak, half rise time, and half decay time were calculated based on average calcium traces (≥3 trials per cell) corresponding to different AP numbers. Peak amplitude was calculated as the difference between the peak identified between -0.02 to 0.28 s to the first AP peak and the baseline as the mean calcium signal between -0.12 to -0.02 s to the first AP peak. Half rise time and half decay time were calculated as the timespans of 0-50% and 100-50% of peak amplitude respectively.

To quantify the sensitivity of calcium indicators, d-prime indices and receiver operating characteristic (ROC) curves were calculated based on ‘signal’ and ‘noise’ of individual trials with ‘signal’ as the peak between -0.02 to 0.28 s to the first AP peak and ‘noise’ the peak between - 0.32 to -0.02 s to the first AP peak. The d-prime index (*d’*) of each cell was calculated using the mean (*μ_signal_*, *μ_noise_*) and variance 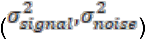 of signal and noise of individual trials (≥3 trials per cell) as:

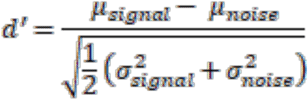

The ROC curve of each AP number was calculated by pooling the signal and noise of individual trials across cells.

### Statistics and reproducibility

Values and errors reported through the text (“### +/- ###”) are mean +/- SEM unless otherwise noted. Error bars and shaded regions are mean +/- SEM. Statistical analyses were performed in GraphPad Prism 8 (in vitro photophysics), and Python (others), as described in the text.

## Author contributions

Conceptualization: A.A., H.A.B., C.D.D., I.-W.C., J.M.G., M.V., T.L., M.W., A.H., R.E.C., and A.W.L.

Data curation: A.A., C.D.D., I.-W.C., P.d.C., J.M.G., J.S.M., M.V., T.L., K.S., R.H.P., C.-Y.W., and K.P.

Formal analysis: A.A., C.D.D., I.-W.C., P.d.C., J.M.G., M.V., T.L., K.S., R.H.P., C.-Y.W., and S.S.

Funding acquisition: C.D.D., R.J.T., T.A.B., Y.N., G.R.J.G., S.M., K.P., A.H., R.E.C., and A.W.L.

Investigation: A.A., C.D.D., I.-W.C., P.d.C., J.M.G., J.S.M., M.V., T.L., K.S., R.H.P., C.-Y.W., and S.S.

Methodology: A.A., H.A.B., C.D.D., I.-W.C., P.d.C., J.M.G., J.S.M., M.V., T.L., K.S., R.H.P., C.- Y.W., F.V., Y.F., and S.S.

Project administration: A.A., G.R.J.G., K.P., A.H., R.E.C., and A.W.L.

Resources: A.A., F.V., Y.F., R.J.T., T.A.B., Y.N., G.R.J.G., S.M., K.P., A.H., R.E.C., and A.W.L. Software: A.A., C.D.D., I.-W.C., M.V., and M.D.N.

Supervision: A.A., M.W., T.T., K.T.-Y., R.J.T., T.A.B., Y.N., G.R.J.G., S.M., K.P., A.H., R.E.C., and A.W.L.

Validation: A.A., C.D.D., I.-W.C., P.d.C., J.M.G., J.S.M., M.V., T.L., K.S., R.H.P., and S.S.

Visualization: A.A., C.D.D., I.-W.C., P.d.C., J.M.G., J.S.M., M.V., T.L., A.H., R.E.C., and A.W.L.

Writing - original draft: A.A., R.E.C., and A.W.L.

Writing - review & editing: A.A., C.D.D., I.-W.C., T.L., K.S., Y.F., T.T., K.T.-Y., G.R.J.G., A.H., R.E.C., and A.W.L.

## Acknowledgments

We thank Cheryl Sank and Karen Ratushny (University of Calgary) for administrative support. We thank Ibis Agosto and Florencia Andres at VectorBuilder for assistance with plasmid orders. We would like to thank Elodie Husi for help with the experiments, and Ronan Chéreau for discussions (University of Geneva). We thank Dr. Rochelin Dalangin for supervising Heather and for discussions.

## References

1. Grienberger, C. & Konnerth, A. Imaging calcium in neurons. Neuron 73, 862–885 (2012).

2. Govorunova, E. G. & Sineshchekov, O. A. The Advances and Applications of Optogenetics. (MDPI, 2020).

3. Rodriguez, E. A. et al. The Growing and Glowing Toolbox of Fluorescent and Photoactive Proteins. Trends Biochem Sci 42, 111–129 (2017).

4. Voigt, F. F. et al. Multiphoton in vivo imaging with a femtosecond semiconductor disk laser. Biomed Opt Express 8, 3213–3231 (2017).

5. Kazemipour, A. et al. Kilohertz frame-rate two-photon tomography. Nat Methods 16, 778– 786 (2019).

6. Kim, D. U. et al. Two-photon microscopy using an Yb(3+)-doped fiber laser with variable pulse widths. Opt Express 20, 12341–12349 (2012).

7. Chen, T.-W. et al. Ultrasensitive fluorescent proteins for imaging neuronal activity. Nature 499, 295–300 (2013).

8. Zhang, Y. et al. Fast and sensitive GCaMP calcium indicators for imaging neural populations. Nature 615, 884–891 (2023).

9. Zhao, Y. et al. An expanded palette of genetically encoded Ca^2+^ indicators. Science 333, 1888–1891 (2011).

10. Dana, H. et al. Sensitive red protein calcium indicators for imaging neural activity. (2016) doi:10.7554/eLife.12727.

11. Shen, Y. et al. A genetically encoded Ca indicator based on circularly permutated sea anemone red fluorescent protein eqFP578. BMC Biol 16, 9 (2018).

12. Katayama, H., Yamamoto, A., Mizushima, N., Yoshimori, T. & Miyawaki, A. GFP-like proteins stably accumulate in lysosomes. Cell Struct Funct 33, 1–12 (2008).

13. Inoue, M. et al. Rational Engineering of XCaMPs, a Multicolor GECI Suite for In Vivo Imaging of Complex Brain Circuit Dynamics. Cell 177, 1346–1360.e24 (2019).

14. Yokoyama, T. et al. A multicolor suite for deciphering population coding of calcium and cAMP in vivo. Nat Methods 21, 897–907 (2024).

15. Mohr, M. A. et al. jYCaMP: an optimized calcium indicator for two-photon imaging at fiber laser wavelengths. Nat Methods 17, 694–697 (2020).

16. Fink, R., et al. PinkyCaMP a mScarlet-based calcium sensor with exceptional brightness, photostability, and multiplexing capabilities. bioRxiv (2025) doi:10.1101/2024.12.16.628673.

17. Dalangin, R. et al. Far-red fluorescent genetically encoded calcium ion indicators. Nat Commun 16, 3318 (2025).

18. Shemetov, A. A. et al. A near-infrared genetically encoded calcium indicator for in vivo imaging. Nat Biotechnol 39, 368–377 (2021).

19. Qian, Y. et al. A genetically encoded near-infrared fluorescent calcium ion indicator. Nat Methods 16, 171–174 (2019).

20. Qian, Y. et al. Improved genetically encoded near-infrared fluorescent calcium ion indicators for in vivo imaging. PLoS Biol 18, e3000965 (2020).

21. Campbell, R. E. et al. A monomeric red fluorescent protein. Proc Natl Acad Sci U S A 99, 7877–7882 (2002).

22. Day, R. N. & Davidson, M. W. The Fluorescent Protein Revolution. (CRC Press, 2014).

23. Shaner, N. C. et al. Improving the photostability of bright monomeric orange and red fluorescent proteins. Nat Methods 5, 545–551 (2008).

24. Wu, J. et al. Improved Orange and Red Ca2+ Indicators and Photophysical Considerations for Optogenetic Applications. (2013) doi:10.1021/cn400012b.

25. Tran, O., Hughes, H. J., Carter, T. & Török, K. Development and characterization of novel jGCaMP8f calcium sensor variants with improved kinetics and fluorescence response range. Front Cell Neurosci 17, 1155406 (2023).

26. Aoyagi, M., Arvai, A. S., Tainer, J. A. & Getzoff, E. D. Structural basis for endothelial nitric oxide synthase binding to calmodulin. EMBO J 22, 766–775 (2003).

27. Wardill, T. J. et al. A Neuron-Based Screening Platform for Optimizing Genetically-Encoded Calcium Indicators. PLOS ONE 8, e77728 (2013).

28. Challis, R. C., et al. Publisher Correction: Systemic AAV vectors for widespread and targeted gene delivery in rodents. Nat Protoc 14, 2597 (2019).

29. Chan, K. Y. et al. Engineered AAVs for efficient noninvasive gene delivery to the central and peripheral nervous systems. Nat Neurosci 20, 1172–1179 (2017).

30. Drobizhev, M., Tillo, S., Makarov, N. S., Hughes, T. E. & Rebane, A. Absolute two-photon absorption spectra and two-photon brightness of orange and red fluorescent proteins. J Phys Chem B 113, 855–859 (2009).

31. Pologruto, T. A., Sabatini, B. L. & Svoboda, K. ScanImage: flexible software for operating laser scanning microscopes. Biomed Eng Online 2, 13 (2003).

32. Pachitariu, M., et al. Suite2p: beyond 10,000 neurons with standard two-photon microscopy. bioRxiv 061507 (2017) doi:10.1101/061507.

33. Institoris, A. et al. Astrocytes amplify neurovascular coupling to sustained activation of neocortex in awake mice. Nat Commun 13, 7872 (2022).

34. Vandal, M. et al. Loss of endothelial CD2AP causes sex-dependent cerebrovascular dysfunction. Neuron 113, 876–895.e11 (2025).

